# SPR-measured dissociation kinetics of PROTAC ternary complexes influence target degradation rate

**DOI:** 10.1101/451948

**Authors:** Michael J. Roy, Sandra Winkler, Scott J. Hughes, Claire Whitworth, Michael Galant, William Farnaby, Klaus Rumpel, Alessio Ciulli

## Abstract

Bifunctional degrader molecules, known as proteolysis-targeting chimeras (PROTACs), function by recruiting a target to an E3 ligase, forming a target:PROTAC:ligase ternary complex. Despite the importance of this key intermediate species, no detailed validation of a method to directly determine binding parameters for ternary complex kinetics has been reported, and it remains to be addressed whether tuning the kinetics of PROTAC ternary complexes may be an effective strategy to improve the efficiency of targeted protein degradation. Here, we develop an SPR-based assay to quantify the stability of PROTAC-induced ternary complexes by measuring for the first time the kinetics of their formation and dissociation in vitro using purified proteins. We benchmark our assay using four PROTACs that target the bromodomains (BDs) of BET proteins Brd2, Brd3 and Brd4 to the E3 ligase VHL. We reveal marked differences in ternary complex off-rates for different PROTACs that exhibit either positive or negative cooperativity for ternary complex formation relative to binary binding. The positively cooperative degrader MZ1 forms comparatively stable and long-lived ternary complexes with either Brd4^BD2^ or Brd2^BD2^ and VHL. Equivalent complexes with Brd3^BD2^ are destabilised due to a single amino acid difference (Glu/Gly swap) present in the bromodomain. We observe that this difference in ternary complex dissociative half-life correlates to a greater initial rate of intracellular degradation of Brd2 and Brd4 relative to Brd3. These findings establish a novel assay to measure the kinetics of PROTAC ternary complexes and elucidate the important kinetic parameters that drive effective target degradation.

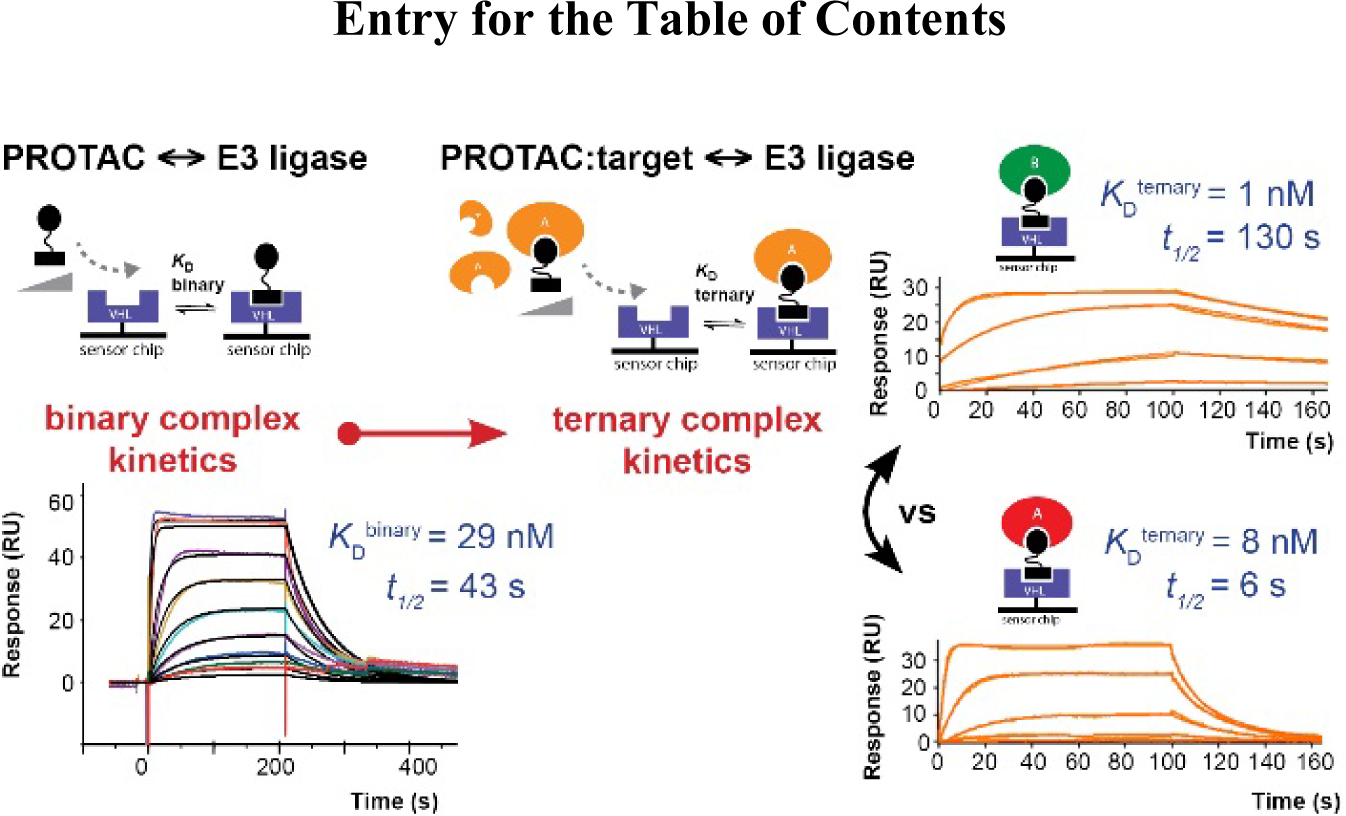

---

PROTACs are bivalent molecules consisting of ligands for each of a target protein and an E3 ligase, joined *via* a linker.*^1, 2^* PROTAC behaviour can be modelled by three-body binding equilibria.*^3^* Formation of a target:PROTAC:ligase ternary complex triggers proximity-dependent target protein ubiquitylation and degradation *via* the ubiquitin-proteasome-system.*^2, 4^* PROTAC drug discovery is rapidly advancing in both academia and industry, fuelled by both improvement in drug-like properties and broader recognition of mechanistic advantages of degradation over inhbition.*^1, 5, 6^* PROTACs offer potential for improved selectivity beyond that of the constituent target ligand, by harnessing additional stabilising or destabilising de novo protein-protein or protein-linker interactions formed *via* the ternary complex.*^2, 7–9^* Thus, in the context of a PROTAC ternary complex (ABC), the binding affinity of the PROTAC ‘B’ to one protein partner ‘C’ (binary binding) may be enhanced or reduced by the presence of the second protein partner ‘A’ (ternary binding). This effect can be quantified in terms of a ‘cooperativity’ (α) factor, defined as the ratio of binary and ternary dissociation constants for PROTAC binding to 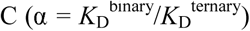 (Figure 1A).*^2^* Cooperativity may be described as ‘positive’ (α > 1, enhanced ternary binding affinity relative to binary, thus further stabilizing the complex), ‘negative’ (α < 1, reduced ternary binding affinity relative to binary, thus destabilizing the complex) or ‘noncooperative’ (α = 1, no change in binding affinity for C due to the presence of A). Developing new tools to understand cooperativity/avidity effects and ternary complex stability in PROTAC design is thus of significant interest. Although functional degraders can be generated in the absence of positive cooperativity,*^8, 10^* mounting evidence suggests enhancing cooperativity and stability of ternary complexes could be an effective strategy in PROTAC design.*^11–13^* As bifunctional molecules, PROTACs are subject to a well-recognised ‘hook effect’, whereby at high PROTAC concentrations, binary interactions may outcompete ternary complex formation.*^5, 143^* Cooperativity is expected to counter this ‘hook effect’ often exhibited by bifunctional molecules, thereby widening the concentration window for PROTAC activity, and could also enable use of weaker-affinity ligands.*^15, 16^*

There is now a growing literature of complimentary methods to measure formation of PROTAC ternary complexes, offering different relative strengths and weaknesses (reviewed more fully elsewhere).*^9^* A number of steady-state methods have been described to measure ternary complex formation and/or cooperativity. These include pull-down assays*^15^* and proximity-based assays (eg. AlphaScreen/AlphaLISA, TR-FRET) to measure relative ternary complex formation at steady state as a function of PROTAC concentration.*^2, 13, 15, 17^* Such assays can offer high throughput, but require labelling of both target and E3 ligase proteins, and the readout of ternary complex formation is indirect and semiquantitative owing to context-specific effects which may potentially affect assay measurement and confound comparison between PROTACs (e.g. effects due to target protein identity, relative protein orientation in the ternary complex, concentrations of proteins used in the assay, intrinsic absorbance or fluorescence of PROTACs). Cooperativity assays based on competitive binding (e.g. fluorescence polarization, to measure PROTAC binding to VHL in the presence or absence of target protein) have also been used to determine binding and cooperativity of PROTACs.*^11^* ITC has also been used for label-free direct quantification of thermodynamic and binding parameters of PROTACs (for both binary and ternary complex formation),*^2, 10, 12^* however is relatively more resource intensive and lower throughput. Notably, none of these assays offer kinetic resolution. Recently, promising cell-based assays have been reported that enable real-time kinetic monitoring of PROTAC ternary complexes in cells, including kinetic EGFP separation of phases-based protein interaction reporter (SPPIER)*^18^* and kinetic bioluminescence resonance energy transfer (BRET)*^19^* approaches. These cell-based methods enable use of full-length proteins, albeit with the caveats that proteins must still be labelled (potentially giving rise to assay-specific effects described earlier) and the readout of ternary complex formation over time is still indirect. Promisingly, the ratiometric nature of BRET enables quantitative measurement.*^19^* However, PROTAC activity in such cell-based assays may also be strongly affected by compound-specific factors including variable cell permeability or non-specific binding to cellular components, as well as the potential general complication of target protein degradation at later time points if normal proteosomal pathway has not been disrupted either genetically or chemically. It is thus also desirable to have well-validated, quantitative biophysical assays to measure PROTAC ternary complex kinetics in vitro using purified proteins.

**Figure 1.**
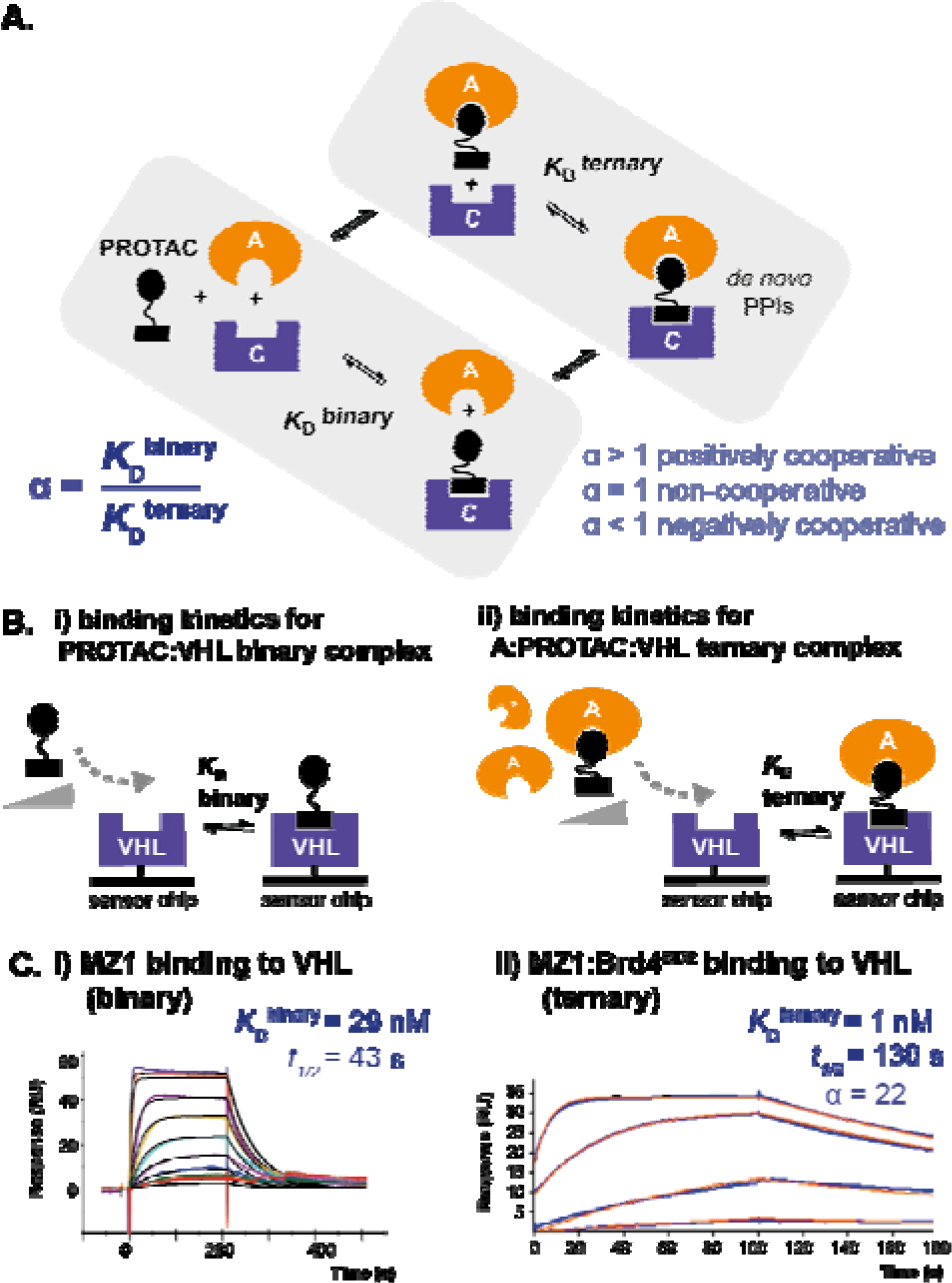
Schematic and binding data illustrating our SPR approach for measuring binding kinetics and determining cooperativity (α) for PROTAC binary and ternary complex formation. A. Ternary binding equilibria, as occur for bivalent molecules (such as PROTACs binding to two proteins, a target ‘A’ and an E3 ligase ‘C’) may involve cooperativity effects, whereby the affinity of the bivalent molecule to one protein (binary complex formation) may be enhanced or decreased when it is already bound to the second protein (ternary complex formation). This may result from additional interactions present in the ternary complex, such as induced *de novo* protein-protein interactions (PPIs). This effect can be represented by a cooperativity factor (α), where 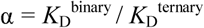. B. To measure the kinetics of PROTAC ternary complexes and determine cooperativity effects, we have developed an SPR assay in which we immobilized the E3 ligase (in our case VHL) onto a sensor chip and measured binding of a PROTAC in either the absence (i) (binary binding) or presence (jj) (ternary binding) of near-saturating concentrations of the target protein (in our case a BET bromodomain). C. Representative SPR binding data are shown using this assay for MZ1 (i) or the MZ1:Brd4^BD2^ complex (ii) binding to immobilised VHL. Binary and ternary binding experiments were performed at 285.15 K in multi-cycle kinetic format and 298.15 K in single-cycle kinetic (SCK) format respectively. For each sensorgram, values shown represent fitted dissociation constants for binary or ternary complex formation (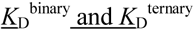 respectively), dissociative half-life of the ternary complex (*t*_1/2_) (calculated as *t*_1/2_ = ln2/*k*_off_) and cooperativity factor (α) (calculated as 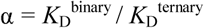).

We were thus interested in exploring surface plasmon resonance (SPR) as a suitable label-free technique to monitor the kinetics of PROTAC-induced ternary complexes, the required intermediate species in the mechanism.*^4^* Herein we develop the first SPR-based assay to quantitatively measure the kinetics of ternary complex formation and dissociation, which we use to characterise the lifetime of ternary complexes composed of bromodomain-containing target proteins, PROTACs, and the von Hippel-Lindau E3 ligase (VHL).

SPR has previously been utilised to characterise three-body binding systems (including complexes composed of protein, DNA and small-molecules),*^20–23^* which can be experimentally involving due to the complex nature of the binding equilibria.*^3^* We sought a general-purpose and conceptually-simple assay format to study PROTACs ternary complexes. Importantly, we recognised for bivalent molecules that the ‘hook effect’ would preclude use of saturating concentrations of PROTAC in the running buffer. We reasoned that by immobilising the E3 ligase, a single sensor surface might be utilised to measure diverse PROTAC/target combinations. To improve uniform presentation on the chip surface, we designed a VHL:ElonginB:ElonginC (VCB) construct harboring an AviTag sequence C-terminal to ElonginB for site-specific biotinylation (hereafter ‘biotin-VHL’).*^24, 25^*

Using a Biacore T200 SPR instrument and streptavidin-immobilised biotin-VHL, we measured the kinetics and affinity of VHL binding for a concentration series of either PROTAC alone (to form binary complex with VHL, 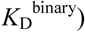 or PROTAC pre-incubated with near-saturating concentrations of target protein (to form ternary complex with VHL, 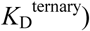 (Figure 1). To achieve near-saturation, the minimum concentration of target protein used was 2 µM (corresponding to approximately 20- to 50-fold in excess of binary *K*_D_ of the PROTAC for the target protein, to ensure at equilibrium ≥ 95-98% formation of binary complex) (refer Supporting Information Figures S1, S2). Experiments were performed in either multi-cycle (binary) or single-cycle (ternary) format without regeneration, to ensure a maximally stable surface and reduce experimental run-times for PROTAC:BD complexes exhibiting slow dissociation kinetics (refer Supporting Information Figure S3).*^26^* Doubly-referenced replicate data were fitted globally (over multiple surface densities where applicable) to a 1:1 Langmuir binding model incorporating a parameter for mass transfer effects, to determine kinetic constants 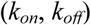 from which dissociation constants were calculated (*K*_D_ = *k*_off_/*k*_on_). Cooperativity (α) was calculated as the ratio 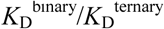.

To benchmark the assay, we utilised the PROTAC MZ1, which forms a highly cooperative ternary complex with VHL and Brd4^BD2^ that we have previously characterised both biophysically and structurally.*^2, 7, 10^* In addition to MZ1, we selected three other PROTACs with ITC-measured affinities for ternary complex formation with VHL/BET bromodomains (Chart 1).*^2, 10^* Combined, these encompass a range of both binary binding affinities and ternary complex cooperativities (both positive and negative). This set includes MZ1 and AT1, two positively-cooperative PROTACs based on a triazolodiazepine BET inhibitor JQ1 (Ligand A, Chart 1), as well as MZP55 and MZP61, two negatively-cooperative PROTACs based on a more potent tetrahydroisoquinolone BET inhibitor I-BET726 (ligand B, Chart 1)

**Chart 1.**
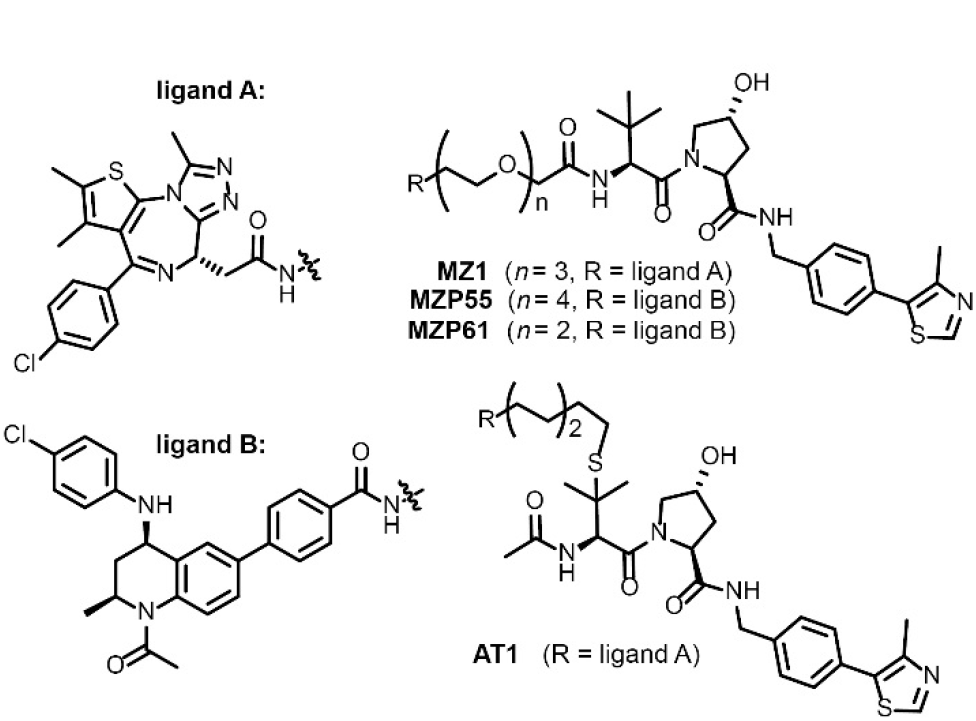
PROTACs utilised in this study.

SPR binding studies were performed with MZ1, AT1, MZP55 and MZP61 alone (binary) or in complex with Brd4^BD2^ as a representative BET bromodomain (ternary), and binding data compared to the ITC-obtained values (Table 1, Supporting Information Table S1, Figures S1 to S4).*^2, 10^* For MZ1, ternary complex formation was also measured with other BET bromodomains (Table 1, Figure 2). No interaction was observed between VHL and Brd4^BD2^ in the absence of PROTAC (Supporting Information Figure S2). For the panel of PROTACs evaluated, the measured *K*_D_ values for binary and ternary complexes, as well as the calculated values for the both cooperativity and change in complex stability (ΔΔG), were remarkably comparable using either ITC or SPR (Table 1). For MZP61 and MZP55 some nonspecific effects were observed for binary binding to VHL (Supporting Information Figure S4); in the case of MZP61 these effects were sufficiently pronounced as to preclude accurate kinetic fitting, so steady state fitting was used to estimate 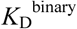.

**Figure 2.**
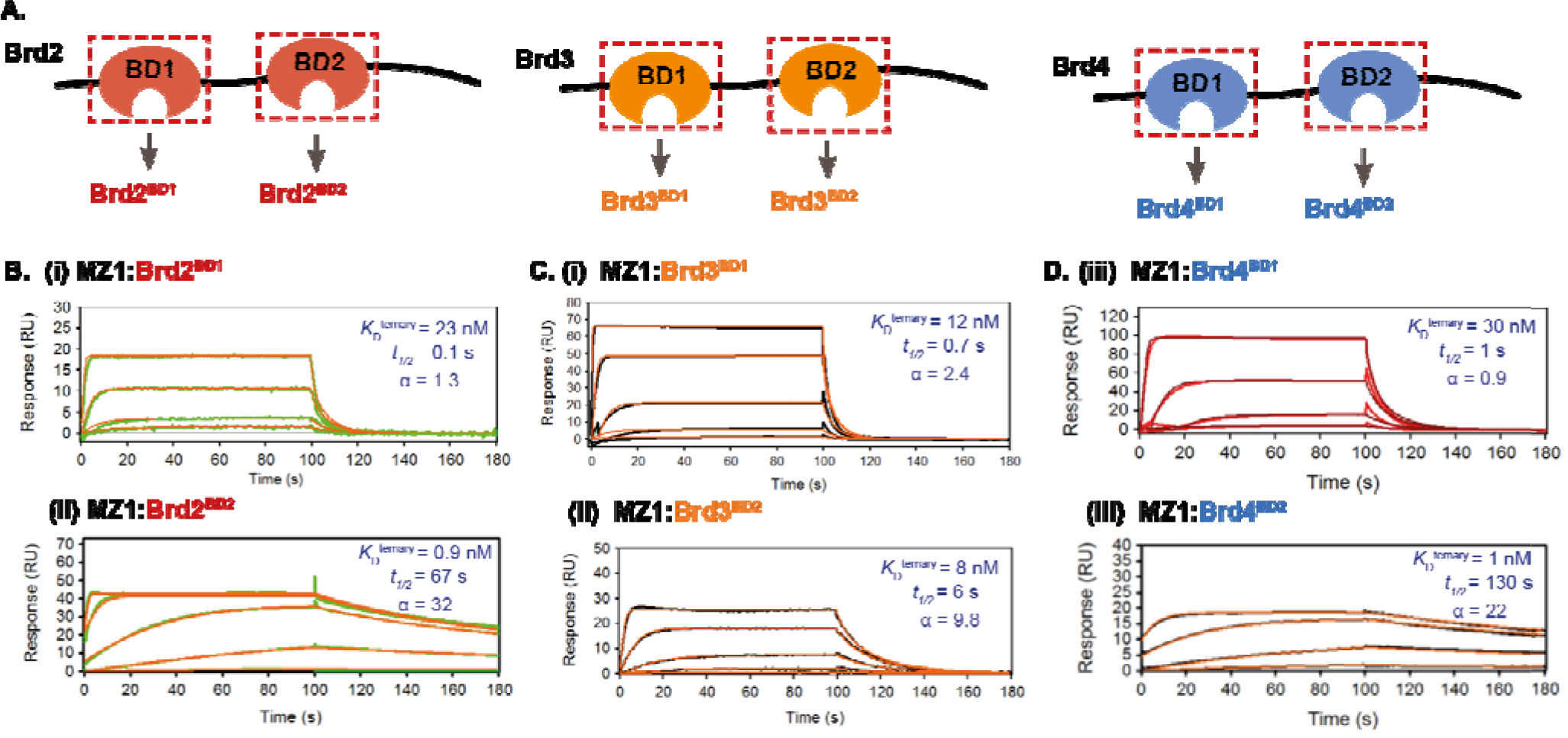
A. Diagram of BET proteins Brd2, Brd3 and Brd4, showing individual bromodomains (BDs) used in this study; these BET proteins exhibit different degradation profiles in response to MZ1 treatment.*^2, 7^* To study the relative kinetics and cooperativity for ternary complex formation with MZ1 and VHL, MZ1:BD complexes were prepared and binding to immobilised VHL measured by SPR. B. SPR sensorgrams for different MZ1:BD complexes reveal marked differences in binding kinetics, in particular VHL:MZ1:Brd2^BD2^ and VHL:MZ1:Brd4^BD2^ ternary complexes dissociated relatively slowly (as a result of the high positive cooperativity, α, and greater complex stability). Ternary binding experiments were performed at 298.15 K in single-cycle kinetic (SCK) format. For each sensorgram, values shown represent fitted dissociation constants for ternary complex formation 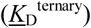, dissociative half-life of the ternary complex (*t*_1/2_) (calculated as *t*_1/2_ = ln2 / *k*_off_) and cooperativity factor (α) (calculated as 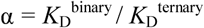).

**Table 1:**
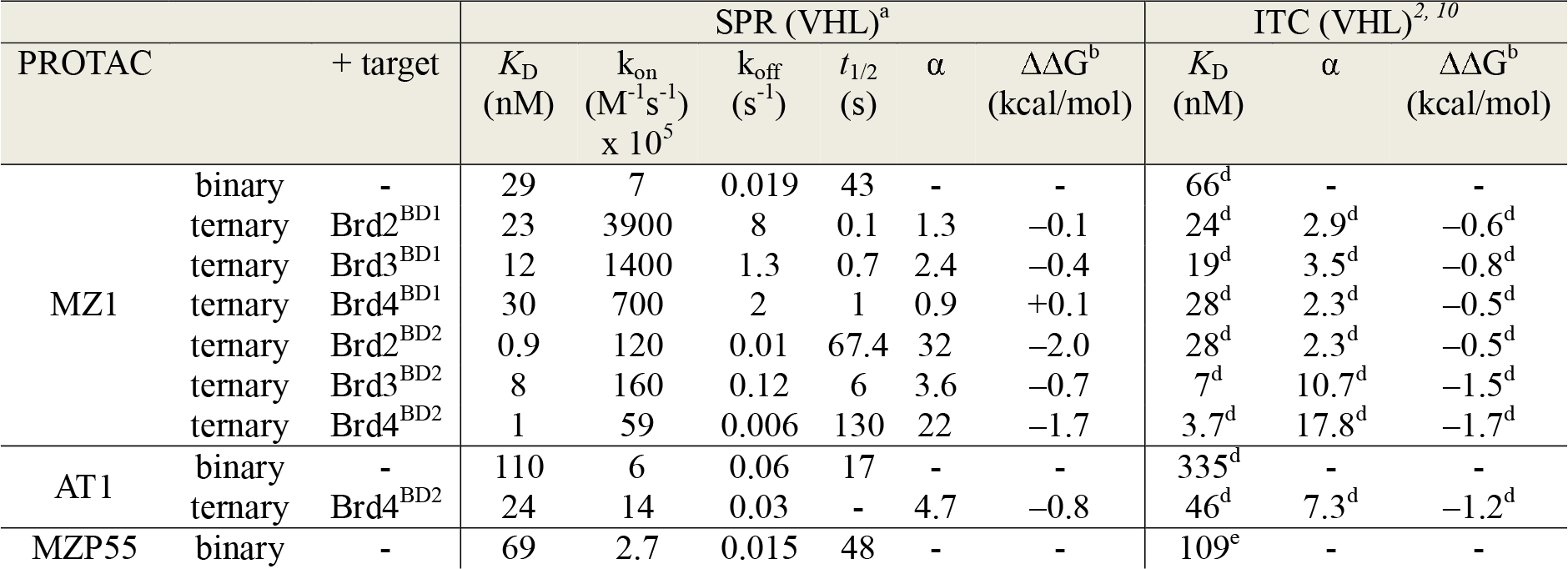
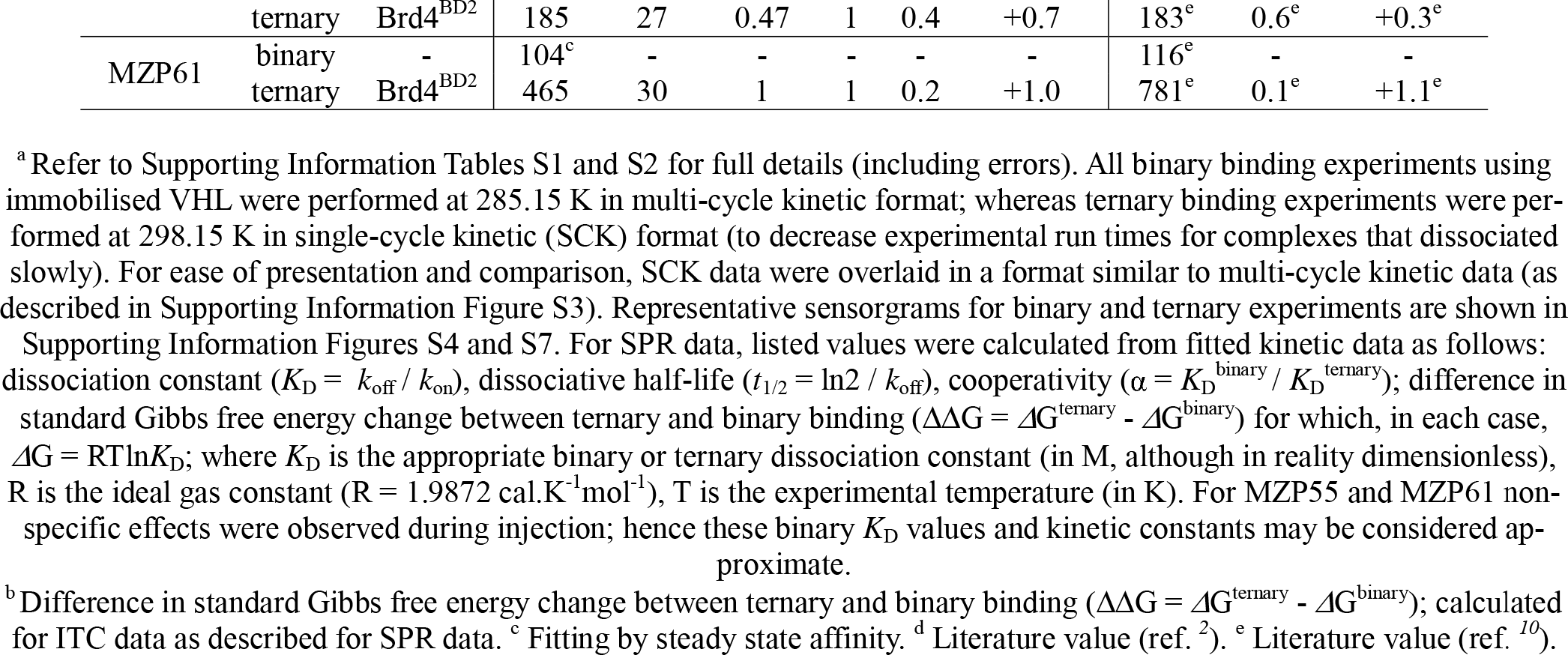
Binding of PROTACs or PROTAC:BD complexes to immobilized VHL (SPR) and comparison to ITC data.

Relative to the binary equilibria, the VHL:MZ1:Brd4^BD2^ ternary complex displayed both a faster *k_on_* and slower *k*_off_, leading to the tighter 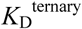 and significant positive cooperativity (α ≈ 20) (Figure 1C, Table 1, Supporting Information Table S1). As anticipated, the VHL:AT1:Brd4^BD2^ ternary complex also exhibited positive cooperativity (α ≈ 5). In stark contrast, ternary complexes formed by either of MZP55 and MZP61 with Brd4^BD2^ had very fast dissociation kinetics (>80-fold faster than the VHL:MZ1:Brd4^BD2^ complex) reflecting overall negative cooperativity, with the ternary complex exhibiting the most negative cooperativity (relative to the binary binding to VHL) being that formed by MZP61 (Supporting Information Table S1, Figure S4). Analogous experiments were conducted in a reversed format, using immobilised Brd4^BD2^ and PROTAC in solution in the absence or presence of excess VHL. Although quantitative fitting of these ternary data was not possible, qualitatively the sensorgrams reflected the same trends already described for the standard format (Supporting Information Figure S6). Whilst we have shown that both the ITC and SPR approaches are complimentary and yield similar values, the notable advantages of our SPR method are increased throughput and yielding kinetic information, including estimates of the lifetime of the ternary complex. This information is invaluable to better understand PROTAC function and ternary complex stability in an enzymatic context,*^4^* analogously to how quantification of inhibitor residence time has helped to understand pharmacological function in certain occupancy-driven small-molecule contexts.*^27, 28^*

MZ1 has been shown to degrade Brd4 more potently than Brd2 or Brd3 (despite near equipotent binding of the constituent warhead ligand to all BET bromodomains),*^7, 10, 19^* resulting from the high thermodynamic stability of the VCB:MZ1:Brd4^BD2^ complex.*^2^* This data suggested varying levels of cooperativity for MZ1 VHL:PROTAC:BD ternary complexes. Moreover, literature data from us using biophysical methods*^2^* and others using cellular degradation assays,*^8^* as well as our own FP data (*vide infra*) are consistent with the MZ1 mediated degradation of BET proteins being driven by complex formation with the BD2s. In an effort to better understand on a kinetic level the basis for apparent differences in cooperativity, we sought to quantify the overall kinetics of VHL:PROTAC:BD ternary complexes for each the different BET bromodomains (Figure 2 and Table 1, Supporting Information Table S2, Figures S7, S8). Ternary complexes consisting of VHL, MZ1 and the first bromodomain (BD1) of either Brd2, Brd3, or Brd4 all displayed very fast dissociation kinetics (t_1/2_ < 1 sec), resulting overall in either low positive cooperativity or no cooperativity (α ≈ 1). In the case of the second bromodomains (BD2), ternary complexes instead exhibited much slower dissociation kinetics (up to 800 slower for a BD2 compared to a BD1 in the case of Brd2, see Table 1), consistent with the known degradation-driving complexes (with BD2s) being the more stable and longer-lived complexes. Crucially, we were struck by the significantly longer ternary half-life of the VCB:MZ1:BD ternary complexes with Brd2^BD2^ and Brd4^BD2^ (t_1/2_≈ 70 and 130 sec respectively) relative to the more short-lived ternary complex with Brd3^BD2^ (t_1/2_ ≈ 6 sec).

Overlay of crystal structures of either Brd2^BD2^ (PDB: 3ONI) or Brd3^BD2^ (PDB: 3S92) in complex with JQ1 and the VCB:MZ1:Brd4^BD2^ structure (PDB: 5T35),*^2^* suggested that ternary complex formation could be influenced by a single amino acid residue difference (Glu^344^ within the ZA loop of Brd3^BD2^, which corresponds to Gly^382^ and Gly^386^ in Brd2^BD2^ and Brd4^BD2^) (Figure 3A). In an equivalent VCB:MZ1:Brd3^BD2^ complex, the side-chain of Glu^344^ would induce severe steric clash with the VHL:MZ1 portion of the complex, leading to destabilisation. We therefore generated reciprocal point mutant swaps (Figure 3B) and measured ternary complex formation by SPR with MZ1 or AT1. In all cases, the resulting SPR binding profiles reflected the effect predicted for the point mutation. The G-to-E point mutation in Brd4^BD2^ or Brd2^BD2^ shortened the ternary complex half-life, decreasing cooperativity and complex stability; whilst the reverse point mutation in Brd3^BD2^ extended the ternary half-life to resemble the profile for Brd4^BD2^ and correspondingly increased cooperativity and stability (Figure 3C, Supporting Information Table S2, Figure S9). As a cross-validation, we evaluated these complexes in a competitive fluorescence polarization (FP) assay; measuring VHL binding of PROTAC or a PROTAC:BD binary complex, *via* displacement of a fluorescent HIF-1α peptide probe (Supporting Information Table S2, Figure S10). Good correlation was observed between SPR-fitted dissociation constants (*K*_D_) and FP-derived inhibition constants (*K*_I_) (Figure 4A). Cooperativity values were also comparable using either method (Supporting Information Table S2). Together, these results underscore the robustness of our SPR approach, and further support the conclusion that the described VCB:MZ1:Brd4^BD2^ structure (PDB: 5T35)*^2^* reflects the predominant (long-lived) species present in solution.

**Figure 3.**
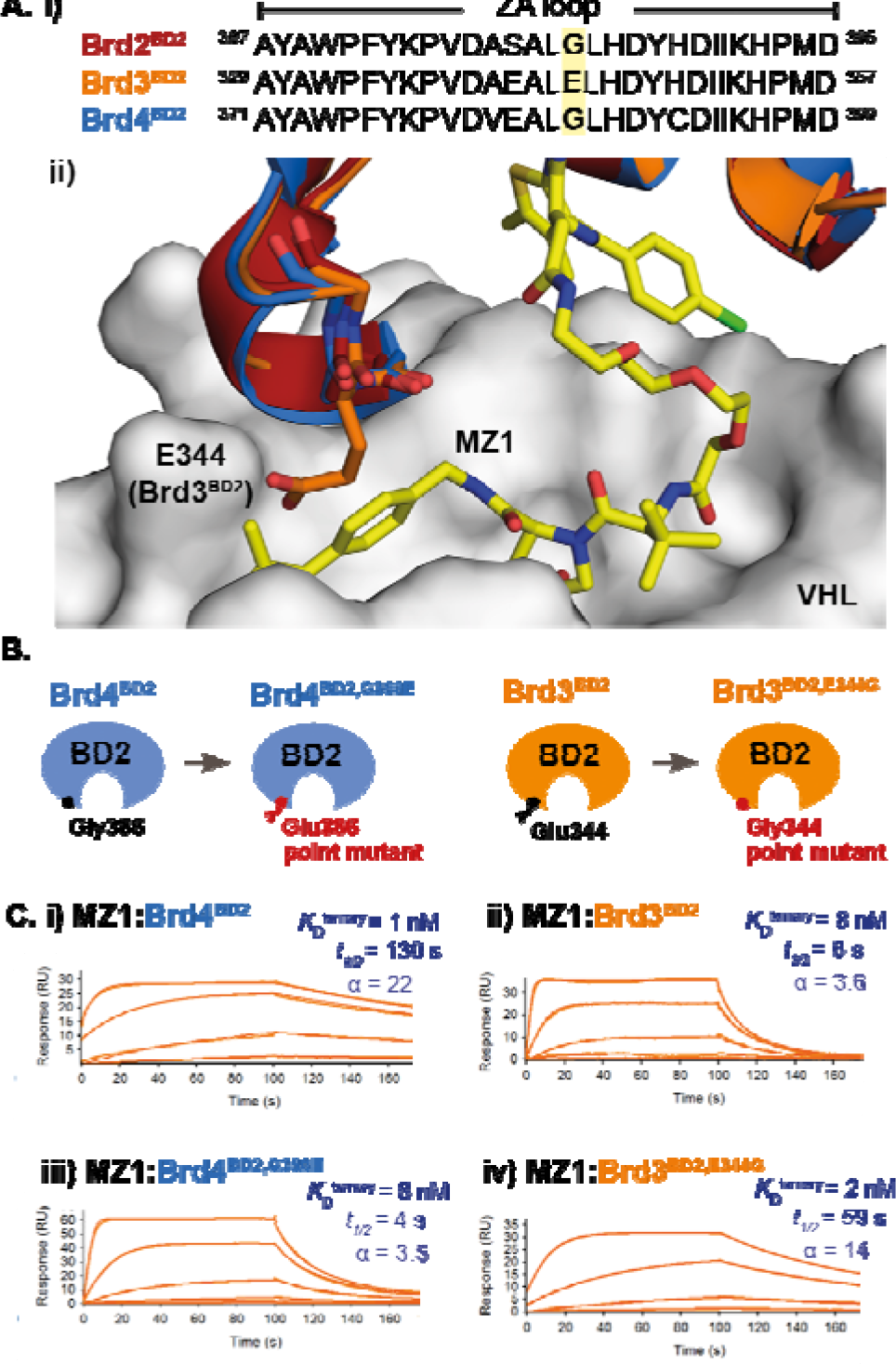
A. Overlay of Brd2^BD2^ (PDB: 3ONI) and Brd3^BD2^ (PDB: 3S92) with the crystal structure of the VCB:MZ1:Brd4^BD2^ ternary complex (PDB: 5T35) suggests that a VHL:MZ1:Brd3^BD2^ ternary complex adopting the equivalent close-packing interaction would likely be less stable due to steric clash with VHL:MZ1 of a single amino acid within the ZA loop of the bromodomain (Glu344 of Brd3^BD2^, which corresponds to Gly386 in Brd4^BD2^). B. Diagram of point mutants generated to explore reciprocal swap of the amino acid at this position. C. Reciprocal exchange of this single Gly/Glu residue in Brd4^BD2^ (i, iii) or Brd3^BD2^ (ii, iv) yields a corresponding swap of the kinetic profile in the resulting VHL:MZ1:BD ternary complex SPR sensorgram. For each sensorgram, values shown represent fitted dissociation constants for ternary complex formation 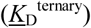, dissociative half-life of the ternary complex (*t*_1/2_) (calculated as *t*_1/2_ = ln2/*k*_off_) and cooperativity factor (α) (calculated as 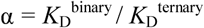).

**Figure 4.**
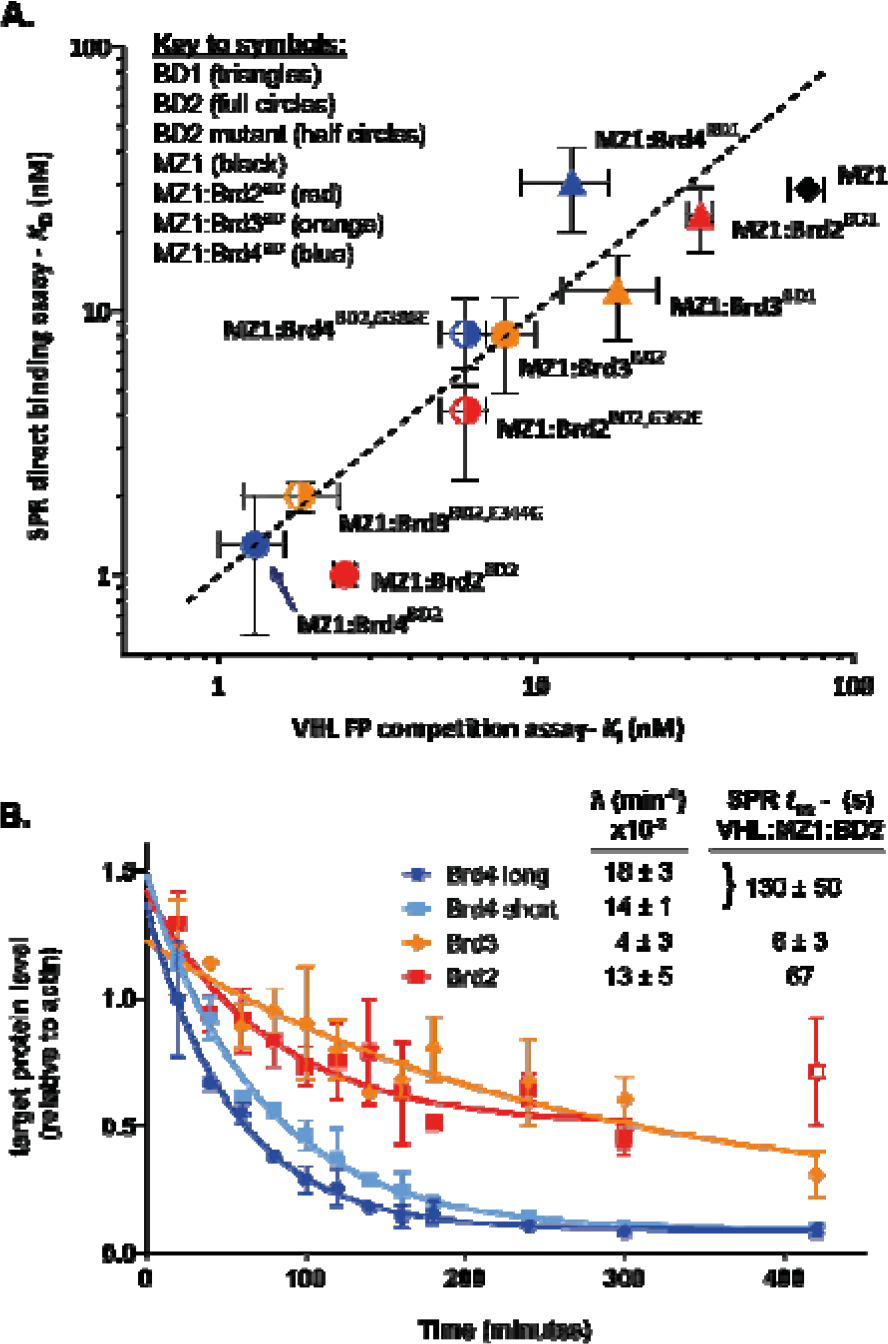
A. Correlation between binary (MZ1) and ternary (MZ1:BD complex) binding to VHL *via* SPR or FP (mean ± SD). B. Initial degradation profile for BET proteins in HEK293 cells in response to MZ1 treatment (333 nM), with initial degradation rates (□) estimated from data fitting (mean ± SEM, N = 3) and SPR ternary half-lives for corresponding VHL:MZ1:BD2 complexes (mean ± SD for N = 2). Note: Short and long isoforms of Brd4 differ in the length of the C-terminus after the tandem bromodomains.*^29^*

These SPR data illustrate kinetically a mechanistic difference between different PROTAC ‘architypes’.*^10^* On the one hand, MZP55 and MZP61 are PROTACs with high binary target affinity (for Brd4), but low or negative cooperativity (Supplementary Information Figures S4 and S6), thus likely forming highly-populated binary complexes but a very transient ternary complex. In contrast, MZ1 and AT1 exhibit weaker binary target affinity (for Brd4), but this is compensated for in the case of Brd4^BD^ by significant positive cooperativity (to form stable ternary complexes). This latter case is predicted to fit a ‘rapid equilibrium’ kinetics model, where a rate-limiting ubiquitination step is dependent on the concentration of PROTAC-induced ternary complex.*^4^* In this regime, an extension in ternary complex stability, and hence lifetime, would be expected to increase rates of target protein ubiquitination and degradation, particularly at early time points prior to countervailing factors such as protein resynthesis or feedback mechanisms. An alternative scenario that might be envisaged for PROTAC molecules operating under a different regime e.g. in a ‘slow-binding regime’, is that their association kinetics might be sufficiently rate limiting as to influence the ultimate outcome of the target degradation.

We wished to examine these hypotheses in a cellular context. Time-course studies were performed to measure initial rate of degradation of Brd4, Brd3, or Brd2 in response to treatment of HEK293 cells with MZ1 (Figure 4B, see Supporting Information Figure S11 for representative western blot data). We observed rapid degradation of Brd2 and both isoforms of Brd4, whilst degradation of Brd3 was significantly slower (Figure 4B, Supporting Information Table S3). Recent results by an independent group, published whilst this manuscript was under preparation, strongly support our conclusion.*^19^* Consistent with our data, Riching *et. al.* observed more rapid initial rates of ubiquitination and degradation of Brd2 and Brd4 than Brd3 in response to MZ1 treatment.*^19^* Crucially, we observed a correlation between the half-life of the VHL:MZ1:BD2 ternary complexes and the initial rate of degradation of the corresponding BET protein, with the more slowly degraded BET protein (Brd3, □ = 4 × 10^3^ min^−1^, cf. with Brd2 and Brd4 which show □ between 13 and 18 × 10^3^ min^−1^) being also the one that exhibits the shortest-lived ternary complex (Figure 4B). This data is consistent with the rapid equilibrium framework, with the more stable and more slowly dissociating ternary complexes driving cellular protein degradation, and not consistent with a slow-binding ternary complex model;*^4^* indeed, MZ1:Brd3^BD2^ showed comparatively fast association kinetics in our SPR data for binding to VHL (k_on_ = 160 × 10^5^ M^−1^s^−1^), possibly even slightly faster as compared to MZ1:Brd2^BD2^ and MZ1:Brd4^BD2^ (Table 1). Taken together, these observations strongly suggest a mechanistic link between the relative half-life of a given target-PROTAC-E3 ligase ternary complex and initial rates of target degradation, which drives a faster and more profound target depletion in cells, at least for PROTACs operating under the ‘rapid equilibrium’ kinetics model such as the archetypical degrader MZ1 studied herein.

In conclusion, we demonstrate a simple and robust SPR-based method to quantify for the first time the stability of target-PROTAC-ligase ternary complexes by measuring the kinetics of their formation and dissociation. We demonstrate that our surface-based SPR method yields values for affinity, cooperativity (α) and complex stability comparable to ITC in solution,*^2, 10^* with increased throughput and yielding additional kinetic information not achievable using other assays described to date. We show by SPR that a single residue can impart significant changes in cooperativity, stability and dissociative half-lives of ternary complexes formed with different but highly conserved target proteins. Lastly, we observe that these kinetic differences of ternary complexes correlate to relative initial rates of target degradation for the well-characterized BET degrader MZ1. Together, these findings establish a new assay for PROTAC ternary complex kinetics and illuminate on ternary complex stability and dissociative half-lives as key optimization parameters for PROTAC design and discovery campaigns.

Our studies suggest a potential challenge/limitation of this approach is the requirement for moderate quantities of whichever target protein is used in ‘near-saturating’ concentrations; however, our experiments also suggest that meaningful data may still be obtained using lower concentrations than that generally used in this study (Supporting Information Figures S1, S5). Our data also suggest that it may be necessary to evaluate both binding orientations (immobilising either the target protein or E3 ligase on the sensor surface), to ascertain which format yields the best data. Nonspecific effects exhibited by certain PROTACs may also pose challenges in determining binary dissociation constants. Notwithstanding this, we anticipate that our SPR kinetic assay will become an established tool to drive PROTAC development and to further elucidate dynamic processes governing their mode of action. We envision one natural extension of our approach may be adaptation to use in more high-throughput assays, in particular due to the additional kinetic information it provides relative to alternative binding approaches such as AlphaScreen or TR-FRET. This is rendered more feasible by the availability of newer highly parallel SPR instruments with improved sensitivity, reducing sample consumption. In our described approach the E3 ligase (in this case VHL) is immobilised on the sensor surface, which has the advantage of being agnostic of the target protein of interest – using this format, any target protein (domain or full-length) might conceivably be used that is capable of recombinant production. In suitable cases, a reversed format using immobilised target protein might provide certain benefits, such as enabling the same PROTAC/E3 ligase solution to be simultaneously screened against both targets and anti-targets on parallel flowcells. Another possibility might be direct capture of over-expressed full-length protein from cell lysate onto the sensor surface by way of a suitable affinity tag. Further, immobilisation of the protein of interest may make this approach capable of extension to screening for the most suitable E3 ligase:PROTAC complexes for forming stable and long-lived complexes. Beyond PROTACs, this assay could also be applied more broadly to study three-body binding equilibria induced by other classes of heterobivalent molecules.*^30^*

## ASSOCIATED CONTENT

### Supporting Information

Supporting results (Tables S1–S3 and Figures S1–S11); supporting materials and methods sections; supporting references.

### Funding Sources

This project has received funding from the European Research Council (ERC) under the European Union’s Seventh Framework Programme (FP7/2007−2013) as a Starting Grant to A.C. (grant agreement no. ERC-2012-StG-311460 DrugE3CRLs), and by Boehringer Ingelheim. Biophysics and drug discovery activities at Dundee were supported by Wellcome Trust strategic awards (100476/Z/12/Z and 094090/Z/10/Z, respectively) to the Division of Biological Chemistry and Drug Discovery.

### Notes

The A.C. laboratory receives sponsored research support from Boehringer Ingelheim and Nurix, Inc. A.C. is a scientific founder, director and shareholder of Amphista Therapeutics, a company that is developing targeted protein degradation therapeutic platforms.

A pre-print of this article (doi: 10.1101/451948) has posted on bioRxiv: http://biorxiv.org/cgi/content/short/451948v1

## ACKNOWLEDGMENT

We thank chemists of the Ciulli group and at B.I. for the gifts of compounds; K. H. Chan for discussions; G. Glendinning and S. Mayer for compound logistics; the Dundee MRC PPU Reagents and Services facility (mrcppureagents.dundee.ac.uk) for the gift of the GST-BirA construct and for DNA sequencing services; and the Dundee Fingerprints Proteomics Facility (proteomics.lifesci.dundee.ac.uk) for ESI-MS of protein samples.

