## Supporting Information for "SPR-measured dissociation kinetics of PROTAC ternary complexes influence target degradation rate"

**Table of Contents:**

Supporting Tables

Table S1. Fitted SPR data for PROTACs (binary) and PROTAC: Brd4$^{BD2}$ complexes (ternary) binding to immobilised VHL and comparison to literature ITC data. **SI3**

Table S2. SPR and FP binding studies with isolated recombinant BET BDs and BET BD point mutants. **SI5**

Table S3. Fitted degradation time course data for HEK293 cells upon treatment with MZ1 (333 nM). **SI7**

Supporting Figures

Figure S1. Selection of PROTAC:target ratio for ternary binding experiments. **SI8**

Figure S2. No significant interaction between Brd4$^{BD2}$ and biotin-VHL is observed in the absence of PROTAC. **SI10**

Figure S3. Illustration of data treatment for ternary single-cycle kinetic (SCK) binding experiments using immobilised biotin-VHL. **SI11 - SI13**

Figure S4. Representative SPR sensorgrams for PROTAC (binary) or PROTAC:Brd4$^{BD2}$ (ternary) binding to immobilised VHL (for PROTACs MZ1, AT1, MZP55, MZP61). **SI14 - SI18**

Figure S5. Effect of varying the PROTAC:target ratio (MZ1:Brd4$^{BD2}$). **SI19 - SI22**

Figure S6. Reversed-format SPR binding experiments (immobilised Brd4$^{BD2}$). **SI23 - SI26**

Figure S7. Representative SPR sensorgrams for MZ1:BD (ternary) binding to immobilised VHL (varying the individual BET bromodomain). **SI27 -SI29**

Figure S8. Representative SPR sensorgrams for AT1:BD (ternary) binding to immobilised VHL (varying the individual BET bromodomain). **SI30 - SI32**

Figure S9. Representative SPR sensorgrams for PROTAC:BD (ternary) binding to immobilised VHL (for PROTACs MZ1 and AT1, and different BET bromodomain point-mutants). **SI33 - SI35**

Figure S10. Fitted Fluorescence Polarization (FP) competition data for MZ1 (binary) and MZ1:BD (ternary) binding to VHL in solution (for individual BET bromodomains and bromodomain point mutants). **SI36**

Figure S11. Representative Western blot for degradation time course data using HEK293 cells. **SI37**

Supporting Experimental Methods

1. Constructs, protein expression and purification **SI38 - SI40**
   - BET bromodomains (BDs) and point mutants **SI38**
   - Chemical biotinylation of Brd4$^{BD2}$ **SI38**
   - VHL and biotinylated VHL **SI39 - SI40**
2. SPR binding studies **SI40 - SI43**
   - SPR binding studies (immobilised VHL) **SI40 - SI42**
   - Reversed-format SPR binding studies (immobilised Brd4$^{BD2}$) **SI42 - SI43**
   - SPR data analysis **SI43**
3. Fluorescence polarization (FP) cooperativity assay (VHL binding) **SI43 - SI44**
4. Cell biology and degradation studies **SI44**
   - Cell lines and culture **SI44**
   - Degradation time course assays **SI44**

Supporting References **SI45**

**Supporting Tables:**

**Table S1. Fitted SPR data for PROTACs (binary) and PROTAC: $Brd4^{BD2}$ complexes (ternary) binding to immobilised VHL and comparison to literature ITC data.**

| | | | | SPR direct binding (VHL)[a] | | | | | | | | | | | ITC (VHL) (literature)[b,1,2] | | | | |
|---|---|---|---|---|---|---|---|---|---|---|---|---|---|---|---|---|---|---|---|
| **PROTAC** | | **+ target** | ***Fit*** | **$k_{on}$ ($M^{-1}s^{-1}$) x $10^5$** | **±** | **$k_{off}$ ($s^{-1}$)** | **±** | **$t_{1/2}$ (s)** | **±** | **$K_D$ (nM)** | **±** | **N** | **α** | **ΔΔG[c] (kcal/mol)** | **$K_D$ (nM)** | **±** | **N** | **α** | **ΔΔG[c] (kcal/mol)** |
| **MZ1** | *binary* | **-** | Kin | 7 | 1 | 0.019 | 0.004 | 43 | 6 | **29** | 3 | 7 | - | - | **66**[d] | 6 | 8 | - | |
| | | | SSA | - | - | - | - | - | **-** | **35** | 5 | 7 | | - | | | | | |
| | *ternary* | $Brd4^{BD2}$ | Kin | 59 | 31 | 0.006 | 0.002 | 130 | 50 | **1** | 1 | 2 | **22** | -1.7 | **3.7**[d] | 0.7 | 2 | **17.8**[d] | -1.7 |
| **AT1** | *binary* | **-** | Kin | 6 | 1 | 0.06 | 0.02 | 17 | 4 | **110** | 40 | 4 | **-** | - | **335**[d] | 30 | 7 | **-** | |
| | | | SSA | - | - | - | - | **-** | **-** | **140** | 50 | 4 | **-** | - | | | | | |
| | *ternary* | $Brd4^{BD2}$ | Kin | 14 | 5 | 0.03 | 0.01 | 26 | 12 | **24** | 5 | 2 | **4.7** | -0.8 | **46**[d] | 6 | 2 | **7.3**[d] | -1.2 |
| **MZP55** | *binary* | **-** | Kin | 2.7 | 0.02 | 0.015 | 0.0001 | 48 | **-** | **69** | - | 1 | **-** | - | **109**[e] | 8 | 2 | **-** | |
| | | | SSA | - | - | - | - | **-** | **-** | **188** | 44 | 3 | **-** | - | | | | **-** | |
| | *ternary* | $Brd4^{BD2}$ | Kin | 27 | - | 0.47 | - | 1.5 | **-** | **185** | - | 1 | **0.4** | +0.7 | **183**[e] | 29 | 2 | **0.6**[e] | +0.3 |
| | | | SSA | - | - | - | - | **-** | **-** | **134** | 25 | 2 | **0.5** | +0.5 | | | | | |
| **MZP61** | *binary* | **-** | SSA | - | - | - | - | **-** | **-** | **104** | 20 | 2 | **-** | - | **116**[e] | 24 | 1 | **-** | |
| | *ternary* | $Brd4^{BD2}$ | Kin | 30 | 30 | 1 | 1 | 0.8 | 0.7 | **465** | 1 | 2 | **0.2** | +1.0 | **781**[e] | 60 | 1 | **0.1**[e] | +1.1 |
| | | | SSA | - | - | - | - | **-** | **-** | **248** | 100 | 2 | **0.4** | +0.6 | | | | | |

Errors are SEM ($N \geq 3$) or SD ($N = 2$).

[a] Analysis where possible is using data from kinetic fitting using a 1:1 Langmuir model including a component for mass transfer effects (Kin), or otherwise using steady state affinity fitting (SSA). SPR binding analyses for binary complexes were performed in multi-cycle kinetic mode at 285.15 K; SPR binding analyses for ternary complexes were performed in single-cycle kinetic (SCK) mode at 298.15 K. For SPR data, listed values were calculated from fitted kinetic data as follows: dissociation constant ($K_D = k_{off} / k_{on}$), dissociative half-life ($t_{1/2} = \ln 2/k_{off}$), cooperativity ($\alpha = K_D^{binary}/K_D^{ternary}$). Nonspecific effects were observed for MZP55 and MZP61.

[b] ITC binding analysis was performed at 298.15 K (literature values).

[c] Difference in standard Gibbs free energy of binding for ternary complex relative to binary ($\Delta\Delta G = \Delta G^{ternary} - \Delta G^{binary}$) for which, in each case, $\Delta G = RT\ln K_D$; where $K_D$ is the appropriate binary or ternary dissociation constant (in M, although in reality dimensionless), R is the ideal gas constant (R = 1.9872 cal.$K^{-1}mol^{-1}$), T is the experimental temperature (in K).

[d] Literature value.[1]

[e] Literature value.[2]

**Table S2. SPR and FP binding studies with isolated recombinant BET BDs and BET BD point mutants.**

| | | | SPR direct binding (VHL)[a] | | | | | | | | | | | FP competition (VHL) | | | |
|---|---|---|---|---|---|---|---|---|---|---|---|---|---|---|---|---|---|
| **PROTAC** | | **+ target** | $k_{on}$ ($M^{-1}s^{-1}$) x $10^5$ | ± | $k_{off}$ ($s^{-1}$) | ± | $t_{1/2}$ (s) | ± | $K_D$ (nM) | ± | N | α | ΔΔG[a] (kcal/mol) | $K_I$ (nM) | ± | N | α |
| **MZ1** | *binary* | - | 7 | 1 | 0.019 | 0.004 | 43 | 6 | **29** | 3 | 7 | - | - | **72** | 9 | 3 | - |
| **MZ1** | *ternary* | Brd2$^{BD1}$ | 3900 | 900 | 8 | 4 | 0.1 | 0.1 | **23** | 9 | 2 | **1.3** | -0.1 | **33** | 3 | 3 | **2.2** |
| | | Brd2$^{BD2}$ | 120 | 20 | 0.0100 | 0.0002 | 67.4 | 0.9 | **0.9** | 0.1 | 2 | **32** | -2.0 | **2.5** | 0.2 | 3 | **29** |
| | | Brd2$^{BD2}$G382E | 130 | 60 | 0.03 | 0.03 | 40 | 30 | **4** | 3 | 2 | **6.9** | -1.1 | **6** | 1 | 2 | **12** |
| | | Brd3$^{BD1}$ | 1400 | 1300 | 1.3 | 0.8 | 0.7 | 0.4 | **12** | 6 | 2 | **2.4** | -0.4 | **18** | 6 | 3 | **4** |
| | | Brd3$^{BD2}$ | 160 | 20 | 0.12 | 0.05 | 6 | 3 | **8** | 5 | 2 | **3.6** | -0.7 | **8** | 2 | 3 | **9.8** |
| | | Brd3$^{BD2}$E344G | 60 | 10 | 0.0120 | 0.0001 | 58.7 | 0.8 | **2.0** | 0.4 | 2 | **14** | -1.5 | **1.8** | 0.6 | 2 | **39** |
| | | Brd4$^{BD1}$ | 700 | 400 | 2 | 1 | 1 | 1 | **30** | 10 | 3 | **0.9** | +0.1 | **13** | 4 | 3 | **5.4** |
| | | Brd4$^{BD2}$ | 59 | 31 | 0.006 | 0.002 | 130 | 50 | **1** | 1 | 2 | **22** | -1.7 | **1.3** | 0.3 | 3 | **55** |
| | | Brd4$^{BD2}$G386E | 1500 | 1900 | 0.2 | 0.2 | 4 | 3 | **8** | 4 | 2 | **3.5** | -0.7 | **6** | 1 | 1 | **12** |
| **AT1** | *binary* | - | 6 | 1 | 0.06 | 0.02 | 17 | 4 | **110** | 40 | 4 | **-** | - | - | - | - | - |
| **AT1** | *ternary* | Brd2$^{BD1}$ | 7 | - | 0.3 | - | 3 | - | **402** | - | 1 | **0.3** | +0.9 | - | - | - | - |
| | | Brd2$^{BD2}$ | 15 | - | 0.04 | - | 20 | - | **27** | - | 1 | **4.1** | -0.7 | - | - | - | - |
| | | Brd2$^{BD2}$G382E | 400 | 400 | 5 | 5 | 0.3 | 0.3 | **150** | 50 | 2 | **0.7** | +0.3 | - | - | - | - |
| | | Brd3$^{BD1}$ | 19 | - | 0.3 | - | 3 | - | **133** | - | 1 | **0.8** | +0.2 | - | - | - | - |
| | | Brd3$^{BD2}$ | 17 | - | 0.3 | - | 3 | - | **163** | - | 1 | **0.7** | +0.4 | - | - | - | - |
| | | Brd3$^{BD2}$E344G | 21 | - | 0.07 | - | 10 | - | **33** | - | 1 | **3.4** | -0.6 | - | - | - | - |
| | | Brd4$^{BD1}$ | 16 | - | 0.9 | - | 0.8 | - | **578** | - | 1 | **0.2** | +1.1 | - | - | - | - |
| | | Brd4$^{BD2}$ | 14 | 5 | 0.03 | 0.01 | 26 | 12 | **24** | 5 | 2 | **4.7** | -0.8 | - | - | - | - |
| | | Brd4$^{BD2}$G386E | 136 | - | 2 | - | 0.4 | - | **133** | - | 1 | **0.8** | +0.2 | - | - | - | - |

Errors are SEM (N ≥ 3) or SD (N = 2).

[a] Analysis where possible is using data from kinetic fitting using a 1:1 Langmuir model including a component for mass transfer effects (Kin), or otherwise using steady state affinity fitting (SSA). SPR binding analyses for binary complexes were performed in multi-cycle kinetic mode at 285.15 K; SPR binding analyses for ternary complexes were performed in single-cycle kinetic (SCK) mode at 298.15 K. For SPR data, listed values were calculated from fitted kinetic data as follows: dissociation constant ($K_D = k_{off} / k_{on}$), dissociative half-life ($t_{1/2} = \ln2/k_{off}$), cooperativity ($\alpha = K_D^{binary}/K_D^{ternary}$), difference in standard Gibbs free energy of binding for ternary complex relative to binary ($\Delta\Delta G = \Delta G^{ternary} - \Delta G^{binary}$) for which, in

each case, $\Delta G = RT\ln K_D$; where $K_D$ is the appropriate binary or ternary dissociation constant (in M, although in reality dimensionless), R is the ideal gas constant (R = 1.9872 cal.$K^{-1}mol^{-1}$), T is the experimental temperature (in K).

**Table S3: Fitted degradation time course data for HEK293 cells upon treatment with MZ1 (333 nM).**

| **Protein** | **$\tilde{\lambda}$ ($min^{-1}$)** | **±** | **$y_0$** | **±** | **Plateau** | **±** | **$t_{1/2}$ (min)** | **±** |
|---|---|---|---|---|---|---|---|---|
| Brd4 long | 0.018 | 0.003 | 1.37 | 0.04 | 0.09 | 0.04 | 38 | 6 |
| Brd4 short | 0.014 | 0.001 | 1.48 | 0.03 | 0.09 | 0.03 | 50 | 5 |
| Brd3 | 0.004 | 0.003 | 1.2 | 0.4 | 0.2 | 0.4 | 176 | 125 |
| Brd2 | 0.013 | 0.005 | 1.4 | 0.1 | 0.5 | 0.1 | 54 | 21 |

Errors are SEM, N = 3.

**a) (i) Selection of PROTAC:target ratio for ternary binding experiements**

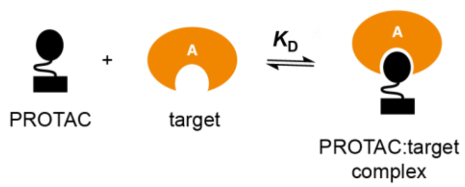


$$\text{fractional occupancy (of PROTAC)} = \frac{[\text{target}]}{[\text{target}] + K_D}$$

**(ii)** To promote >95% formation of PROTAC:target binary complex (at equilibrium in each well prior to injection):

1. [target] > 20-fold over $K_D$
2. [target] > [PROTAC]

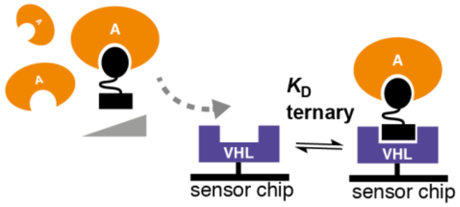


**b) Reported binary affinity of PROTACs for isolated bromodomains**

| | Reported affinity (ITC) $K_D$ (nM) | | | | | |
|---|---|---|---|---|---|---|
| **Compound** | **$BRD2^{BD1}$** | **$BRD2^{BD2}$** | **$BRD3^{BD1}$** | **$BRD3^{BD2}$** | **$BRD4^{BD1}$** | **$BRD4^{BD2}$** |
| **MZ1** | 62 [a] | 60 [a] | 21 [a] | 13 [a] | 39 [a] | 26 [b]<br>15 [a] |
| **AT1** | 111 [a] | 94 [a] | 35 [a] | 39 [a] | 75 [a] | 44 [a] |
| **MZP61** | - | - | - | - | - | 3 [b] |
| **MZP55** | 45 [b] | 5 [b] | 39 [b] | 11 [b] | 39 [b] | 8 [b] |

[a] Literature value (ref. [1])

[b] Literature value (ref. [2])

**Figure S1. Selection of PROTAC:target ratio for ternary binding experiments.** (a) For a reversible 1:1 interaction between a PROTAC and target protein that follows the law of mass action, by analogy to traditional descriptions of receptor-ligand binding,[3] the Hill-Langmuir equation leads to the right-hand equation in (a)(i) which describes the fractional occupancy at equilibrium of the PROTAC by the target protein. This relates the fractional occupancy of the PROTAC to the concentration of the target protein (shown as [target]) and the dissociation constant ($K_D$) of the binary PROTAC-target interaction. Based on this relationship, for ternary binding experiments, we elected to pre-incubate the PROTAC with a near-saturating concentration of the target protein (corresponding to at least 20-fold in excess of the binary $K_D$ of the PROTAC/target protein interaction, and at all times in stoichiometric excess relative to the concentration of PROTAC), to ensure a minimum binary occupancy of greater than 95%. This ensures that the concentration of free PROTAC remaining in the injected well solution, which will also compete with the PROTAC:target binary complex for binding to the immobilised E3 ligase, remains negligible. The binary affinities for the majority of the PROTAC/bromodomain pairs used in this study have previously been determined (listed in (b); shown to be 1:1 interactions as measured by ITC),[1, 2]. Based on these values, for PROTAC ternary binding experiments using immobilised biotin-VHL, we elected to set the minimum concentration of free 'near-saturating' bromodomain in each well solution to be 2 μM; such that, for all PROTACs tested, the fraction of binary PROTAC:BD complex formed prior to injection would be expected to be in the range 95 to 98%. We deemed this concentration an acceptable compromise between consumption of target protein and the expected accuracy of the measured binding response; this decision will almost certainly vary according to the nature of the experiment and the interacting partners. The equivalent calculation can also be made for other types of ternary interaction, or for PROTACs, if the reversed orientation is used (i.e. the target protein immobilised and excess E3 ligase is used in solution).

**a) MZ1:Brd4$^{BD2}$ (ternary) binding to immobilised biotin-VHL**

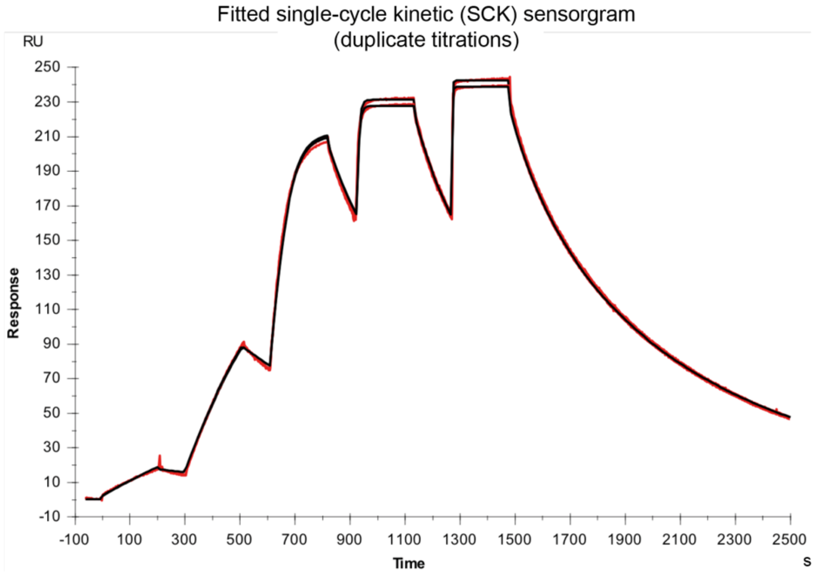


**b)**

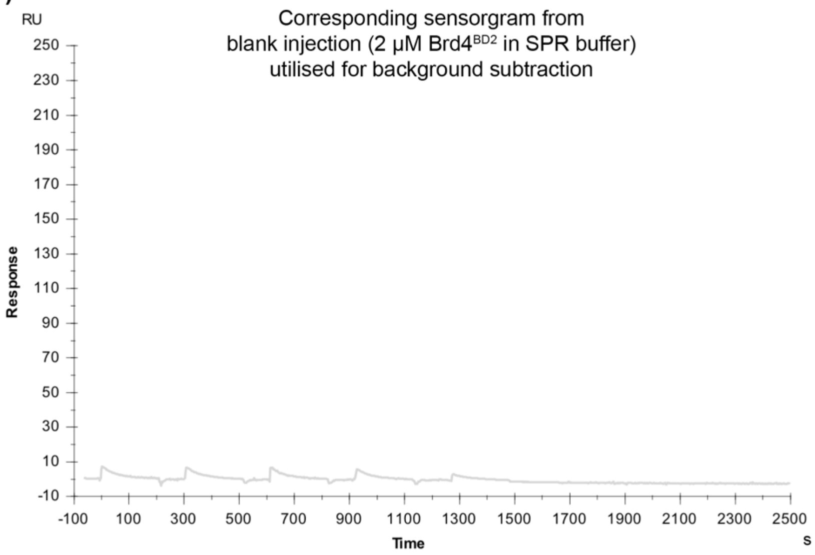


**Figure S2. No significant interaction between Brd4$^{BD2}$ and VHL is observed in the absence of PROTAC.** Sensorgrams are shown for a representative ternary single-cycle kinetic (SCK) experiment to measure binding of MZ1-Brd4$^{BD2}$ to immobilised biotin-VHL. The first sensorgram (a) shows the double-referenced binding data for sequential injection of increasing concentrations MZ1 (1.6 nM to 1000 nM) in the presence of near-saturating concentrations of Brd4$^{BD2}$ (2 to 25 µM) over immobilised biotin-VHL, resulting a binding response due to ternary complex formation in the presence of PROTAC. The second sensorgram (b) is the corresponding binding response from a series of blank injections (2 µM Brd4$^{BD2}$ in SPR buffer) used for background subtraction. No significant binding of Brd4$^{BD2}$ to VHL is observed in the absence of PROTAC; as was also the case for all other purified bromodomains used in this study (data not shown).

**a) MZ1:Brd4$^{BD2}$ (ternary)**
**Binding to immobilised biotin-VHL**

Ternary binding experiments were performed in single cycle kinetic (SCK) format. Shown are overlaid SCK sensorgrams from four replicate titrations (of 5 sequential injections each) over three flowcells immobilised with different surface densities of biotin-VHL.

**(i)**

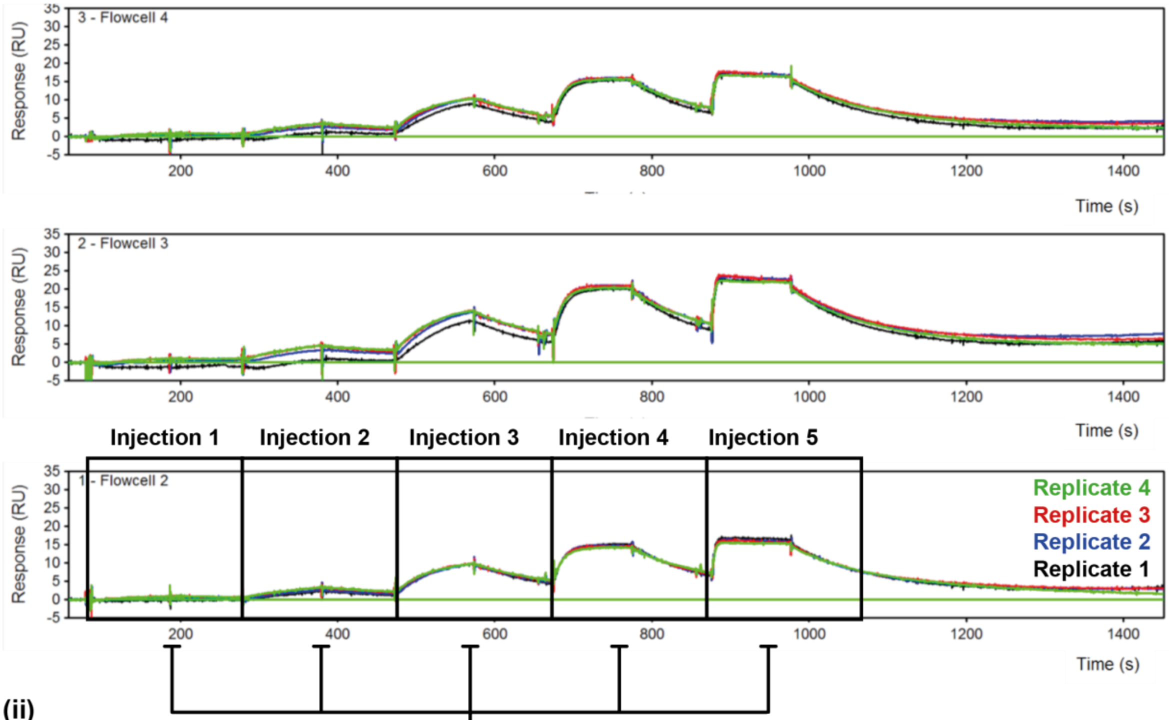


**(ii)**

**Overlay/fitting of injections for each replicate on Flowcell 2.** Representive overlays from a single flowcell are show below.

For each replicate, kinetic constants were obtained by simultaneous global fitting of data from three flowcells (different surface densities of biotin-VHL).

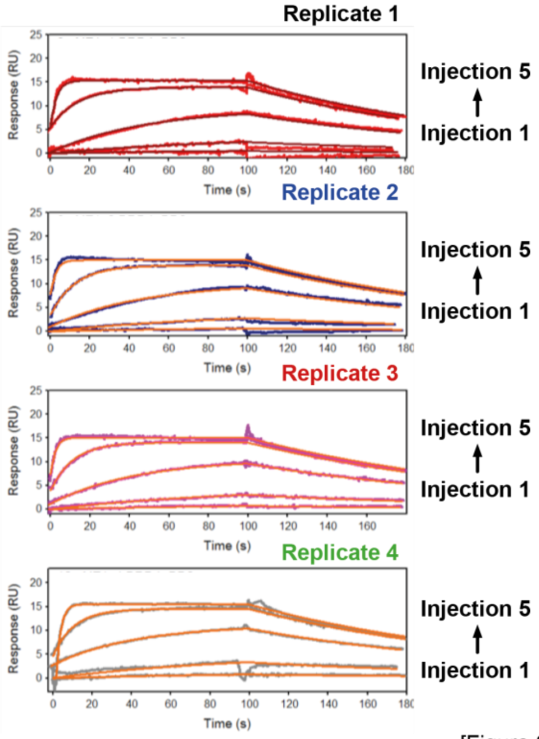


[Figure S3 continues on the next page.]

**b) MZ1:Brd3$^{BD2}$ (ternary)**
**Binding to immobilised biotin-VHL**

Ternary binding experiments were performed in single cycle kinetic (SCK) format. Shown are overlaid SCK sensorgrams from four replicate titrations (of 5 sequential injections each) over three flowcells immobilised with different surface densities of biotin-VHL.

**(i)**

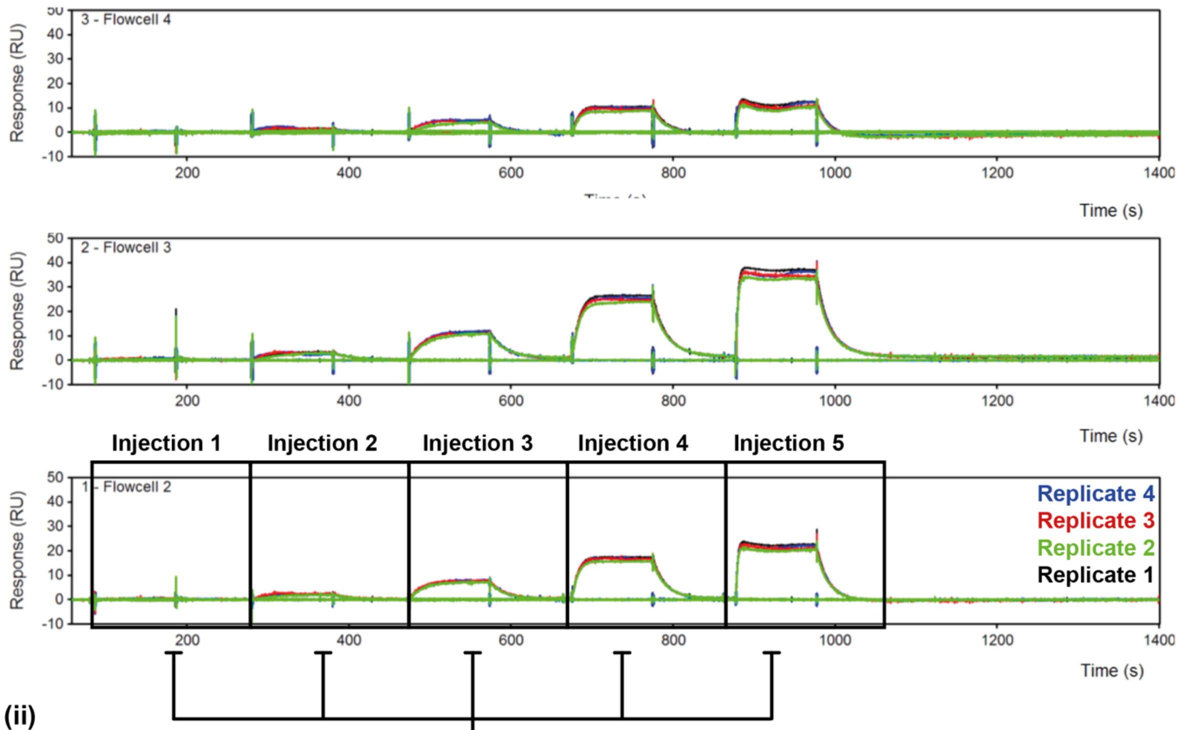


**(ii)**

**Overlay/fitting of injections for each replicate on Flowcell 2.** Representive overlays from a single flowcell are shown below.

For each replicate, kinetic constants were obtained by simultaneous global fitting of data from three flowcells (different surface densities of biotin-VHL).

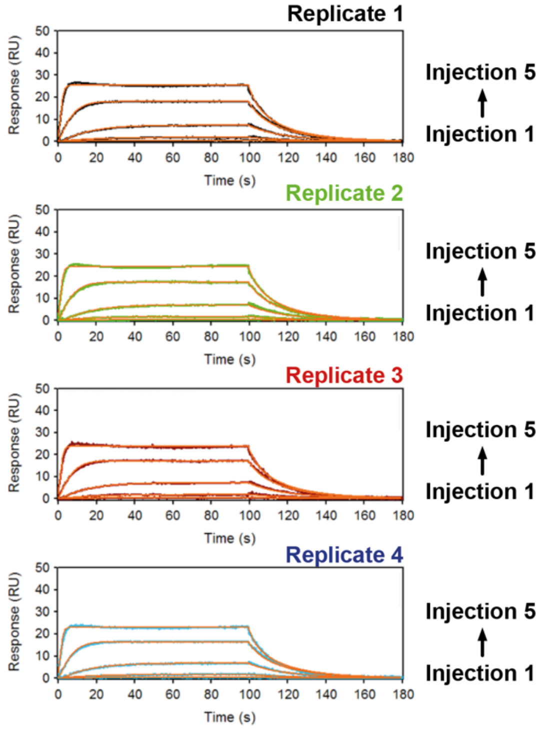

**Figure S3 (preceding page). Illustration of data treatment for ternary single-cycle kinetic (SCK) binding experiments using immobilised biotin-VHL.** Single representative experiments are shown for MZ1:Brd4$^{BD2}$ (a) and MZ1:Brd4$^{BD2}$ (b). For each ternary binding experimental repeat, three to four replicate titrations were performed over two to three flowcells with different immobilised surface densities of biotin-VHL. To facilitate comparison between ternary complexes and with binary multicycle data, as well as to make more efficient use of space in figures, sequential injections for ternary SCK experiments were overlaid in a format similar to that typically used for multi-cycle experiments. This was done for each replicate, by shifting the X-axis to align the injection time, as illustrated above. Double-referenced data were then globally fitted simultaneously over all flow cells using a 1:1 Langmuir interaction model, with a term for mass-transport included (processed using zip fitting in Scrubber) (BioLogic Software).

**Binding to immobilised biotin-VHL:**

**a) MZ1** **Binary**

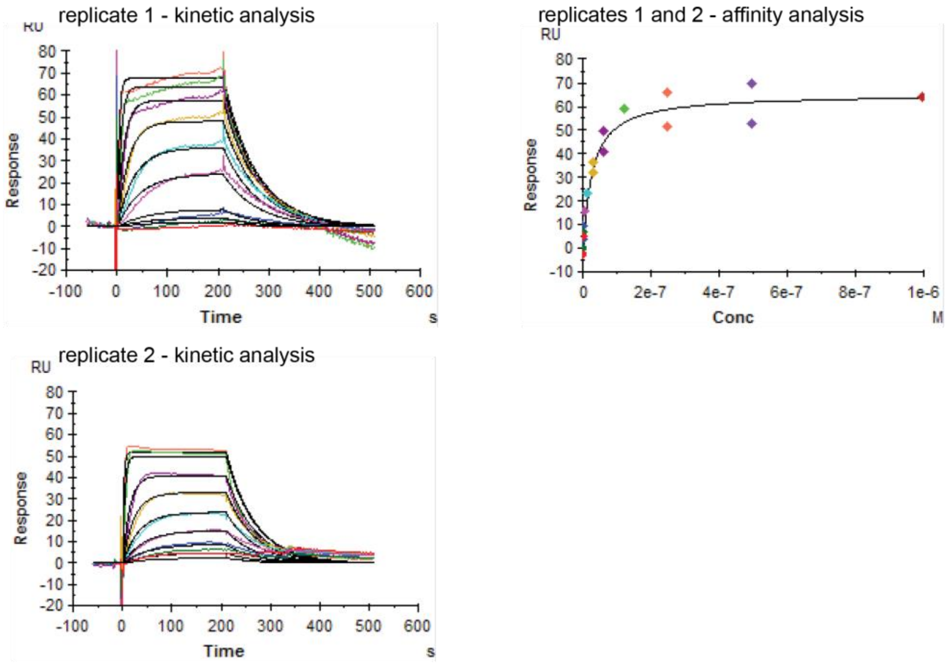


**b) MZ1:Brd4$^{BD2}$** **Ternary**

All samples globally fitted (kinetic analysis)

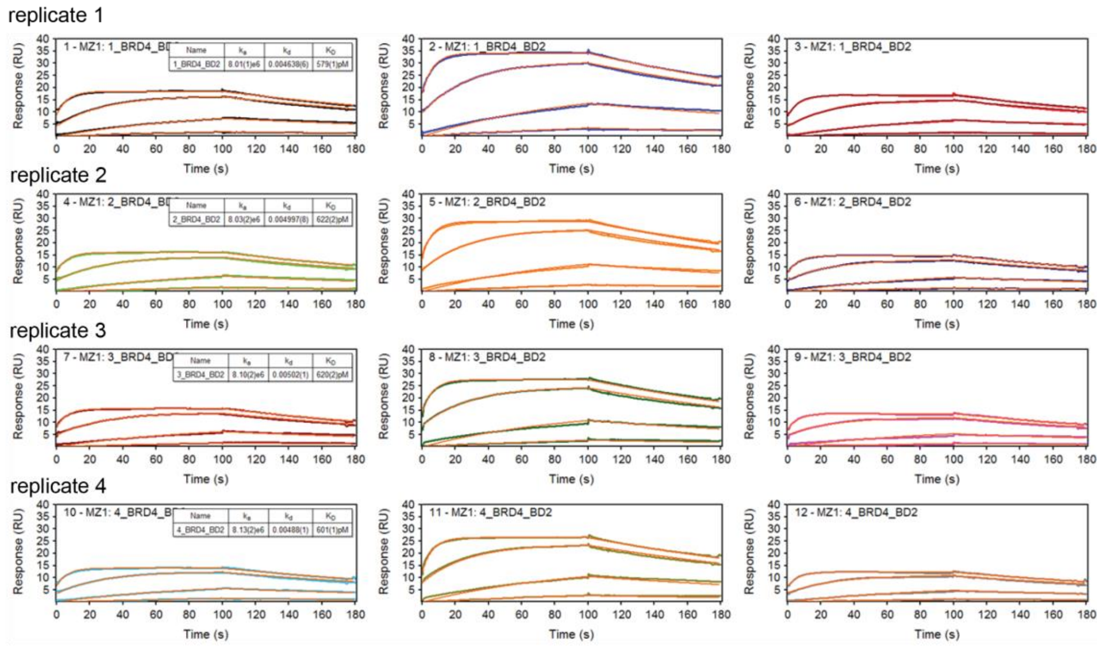


[Figure S4 continues on the next page.]

**Binding to immobilised biotin-VHL:**

**c) AT1** **Binary**

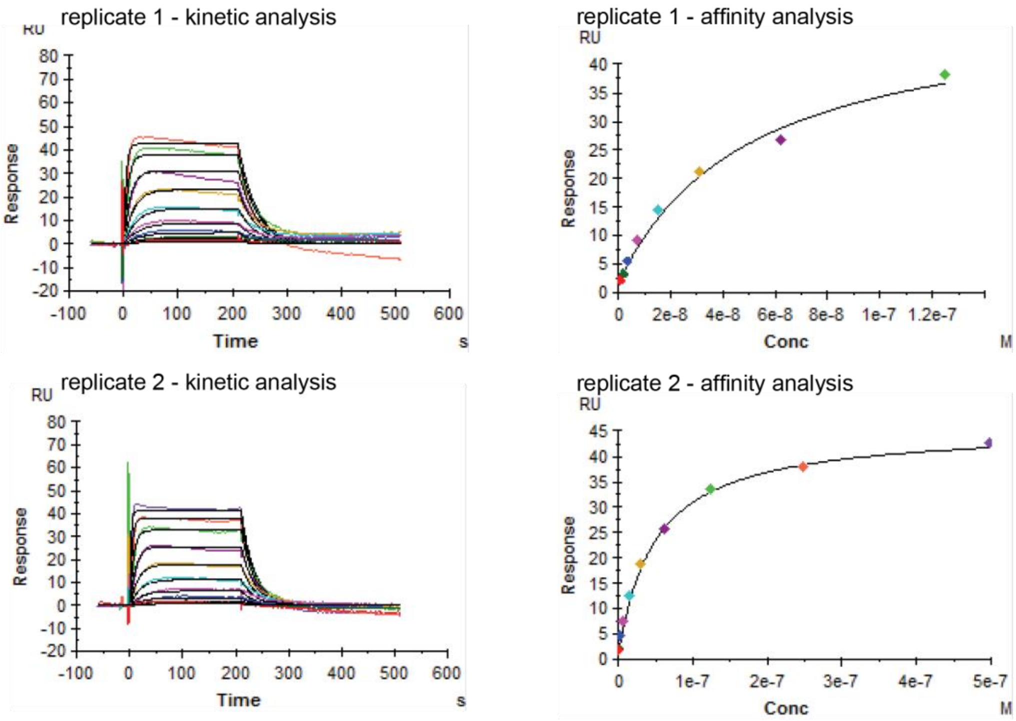


**d) AT1:Brd4[BD2]** **Ternary**

All samples globally fitted (kinetic analysis)

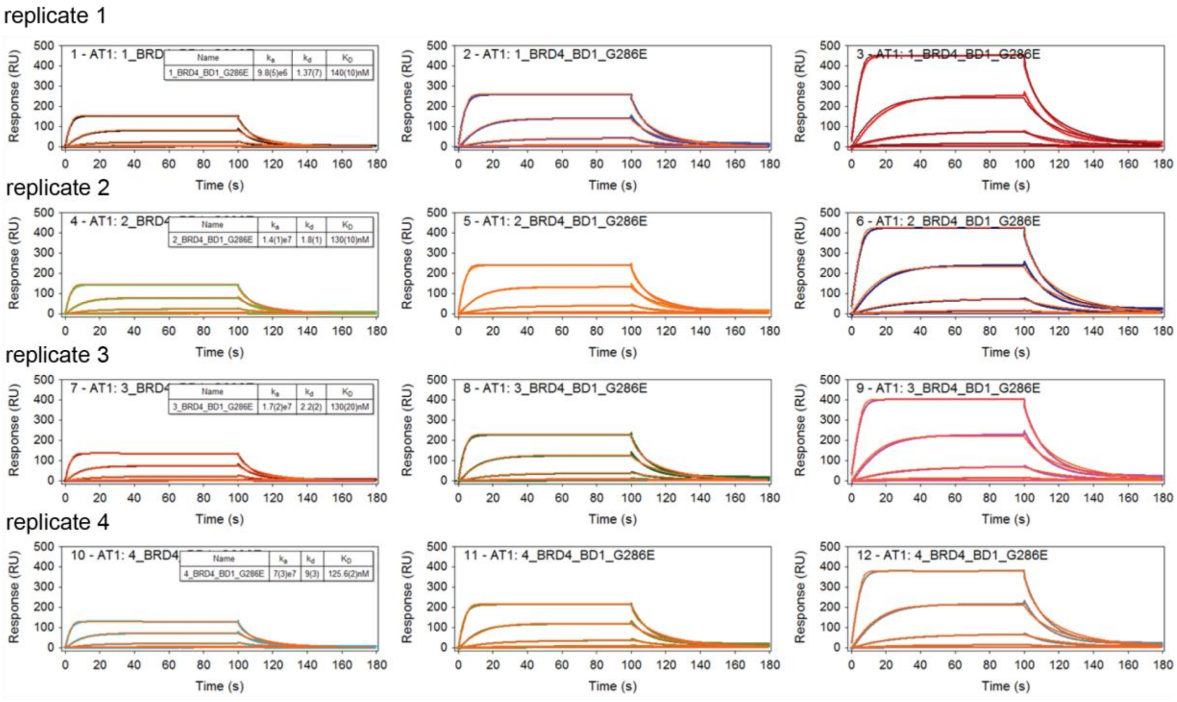


[Figure S4 continues on the next page.]

**Binding to immobilised biotin-VHL:**

**e) MZP55** **Binary**

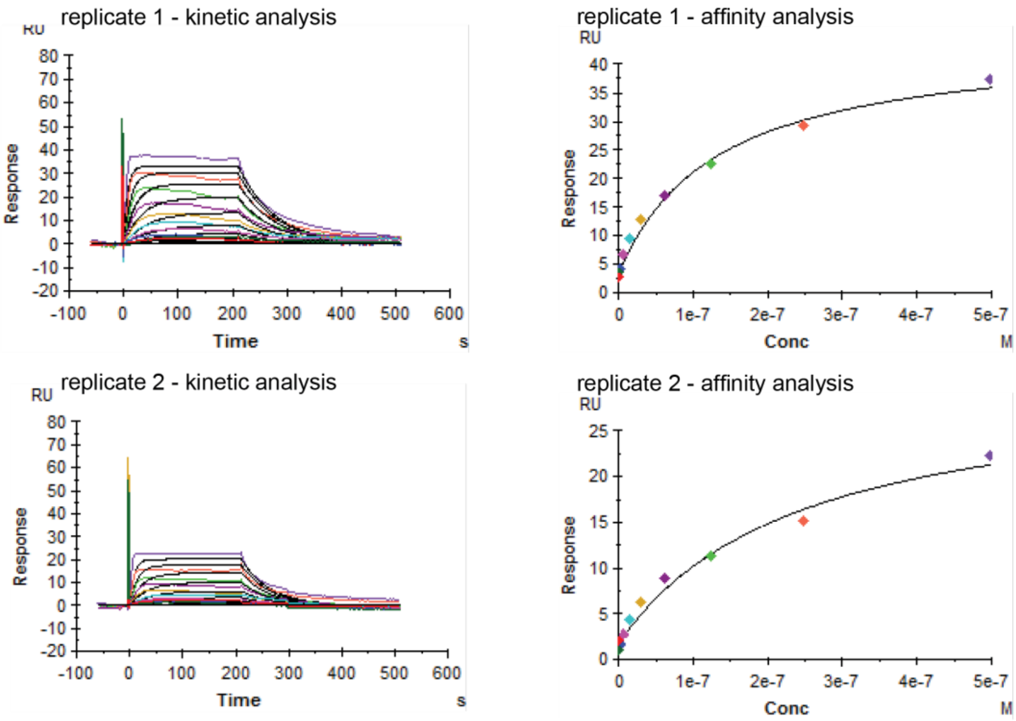


**f) MZP55:Brd4$^{BD2}$** **Ternary**

All samples globally fitted (kinetic analysis)

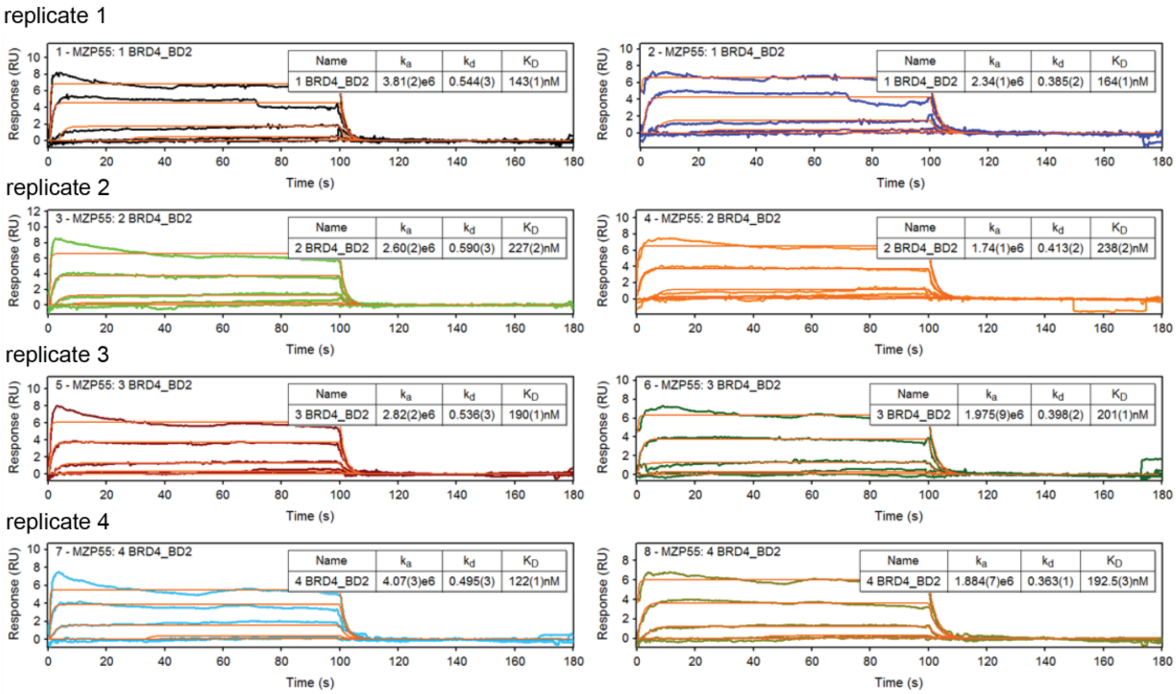


[Figure S4 continues on the next page.]

**Binding to immobilised biotin-VHL:**

**g) MZP61**

**Binary**

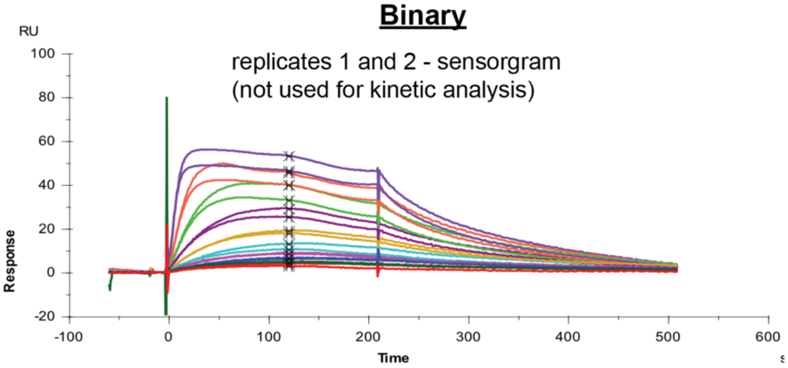


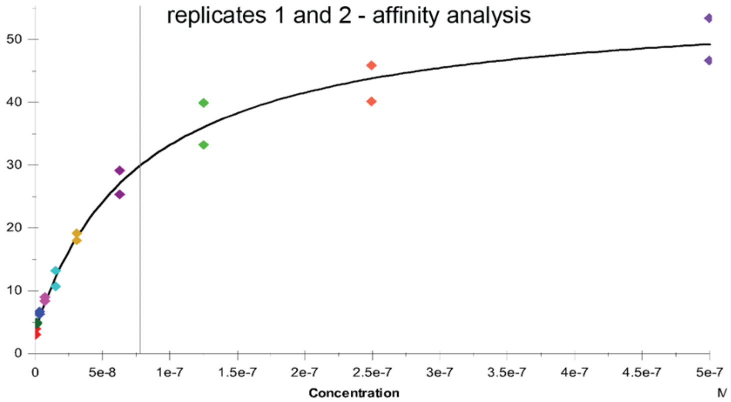


**h) MZP61:Brd4$^{BD2}$**

**Ternary**

All samples globally fitted (kinetic analysis)

replicate 1

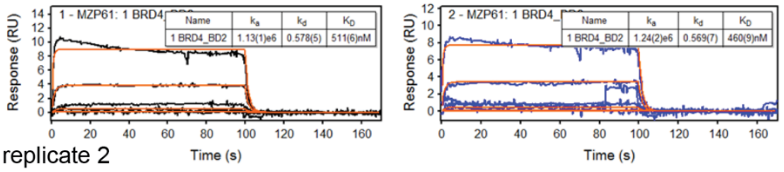


replicate 2

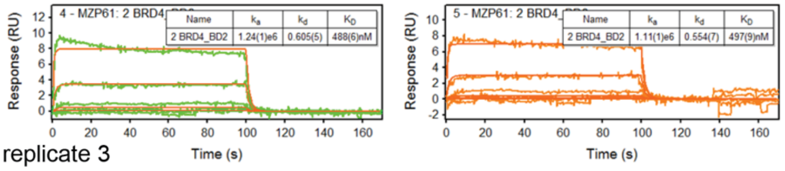


replicate 3

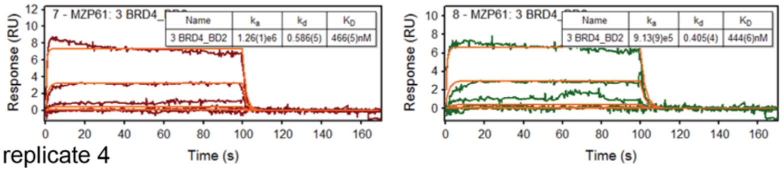


replicate 4

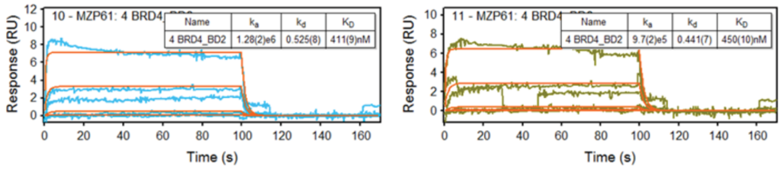

**Figure S4 (preceding page). Representative SPR sensorgrams for PROTAC (binary) or PROTAC:Brd4$^{BD2}$ (ternary) binding to immobilised VHL (for PROTACs MZ1, AT1, MZP55, MZP61).** SPR binding analyses for binary complexes were performed in multi-cycle kinetic mode at 285.15 K; SPR binding analyses for ternary complexes were performed in single-cycle kinetic (SCK) mode at 298.15 K and analysed as described (Figure S3). For MZP55 and MZP61 nonspecific effects were observed during the second half of injection; hence these binary $K_D$ values may be considered estimates. For MZP61, only steady state fitting was performed.

### a) Effect of varying the PROTAC:target ratio (MZ1:Brd4$^{BD2}$)

#### (i) Overview of experiment

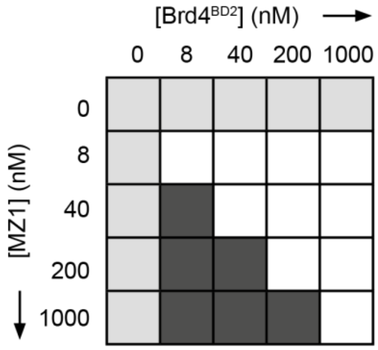


#### (ii) Fitted SPR data at varying (MZ1:Brd4$^{BD2}$) ratios (binding to immobilised biotin-VHL at 285.15 K)

| Ratio (MZ1:Brd4$^{BD2}$) | [MZ1] (nM) | $k_{on}$ (1/Ms) x $10^5$ | $k_{off}$ (1/s) | $K_D$ (nM) | $R_{max}$ (RU) | Chi² (RU²) |
|---|---|---|---|---|---|---|
| 1:1 | 8 | 7.8 | 0.0026 | 3.28 | 88.86 | 0.08 |
| | 40 | 9.4 | 0.0026 | 2.72 | 113.4 | 0.62 |
| | 200 | 6.2 | 0.0024 | 3.94 | 114.9 | 1.12 |
| | 1000 | 3.9 | 0.0029 | 7.26 | 119.2 | 2.28 |
| 1:5 | 8 | 24.0 | 0.0029 | 1.21 | 110.9 | 0.26 |
| | 40 | 26.5 | 0.0030 | 1.12 | 117.2 | 0.83 |
| | 200 | 25.4 | 0.0034 | 1.34 | 119 | 1.67 |
| 1:25 | 8 | 34.4 | 0.0031 | 0.89 | 112.7 | 0.45 |
| | 40 | 38.1 | 0.0037 | 0.98 | 115.9 | 6.61 |
| 1:125 | 8 | 34.5 | 0.0032 | 0.93 | 111.5 | 0.82 |

### b) Fitted SPR binding constants for MZ1:Brd4$^{BD2}$ at varying PROTAC:target ratios

(i)

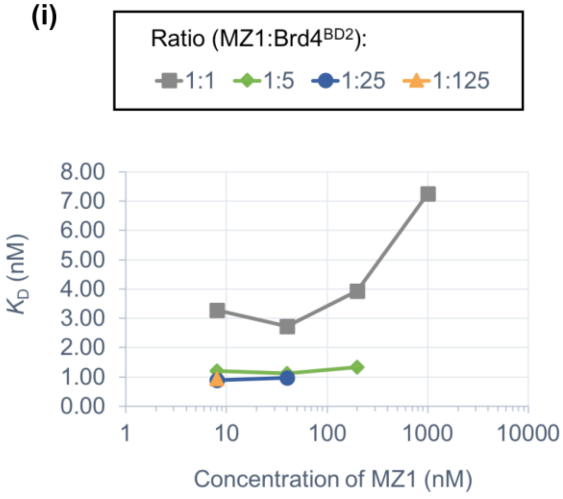


(ii)

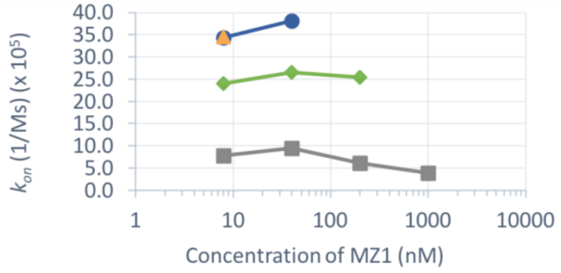


(iii)

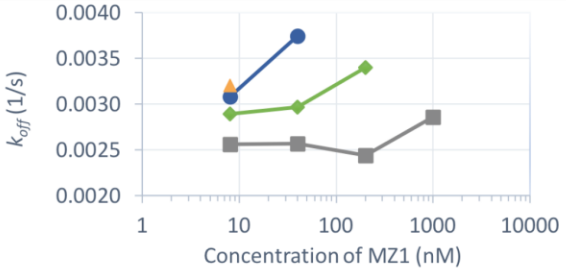


(iv)

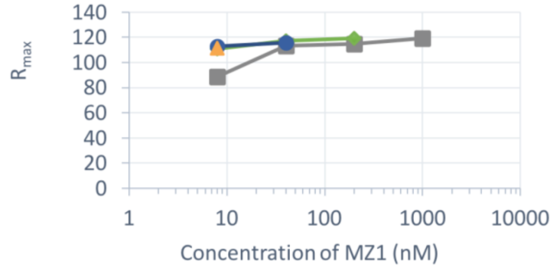


[Figure S5 continues on the next page.]

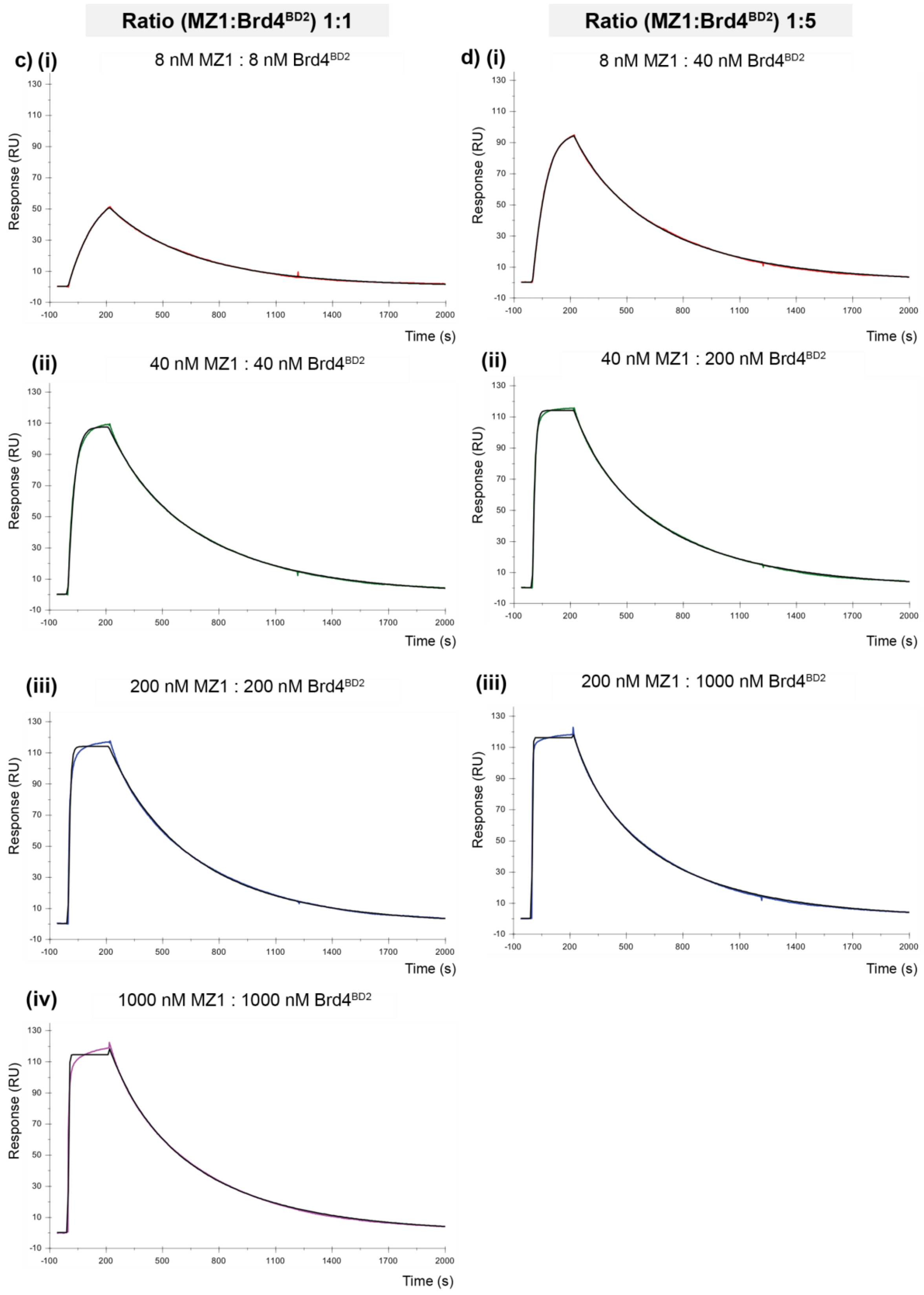


[Figure S5 continues on the next page.]

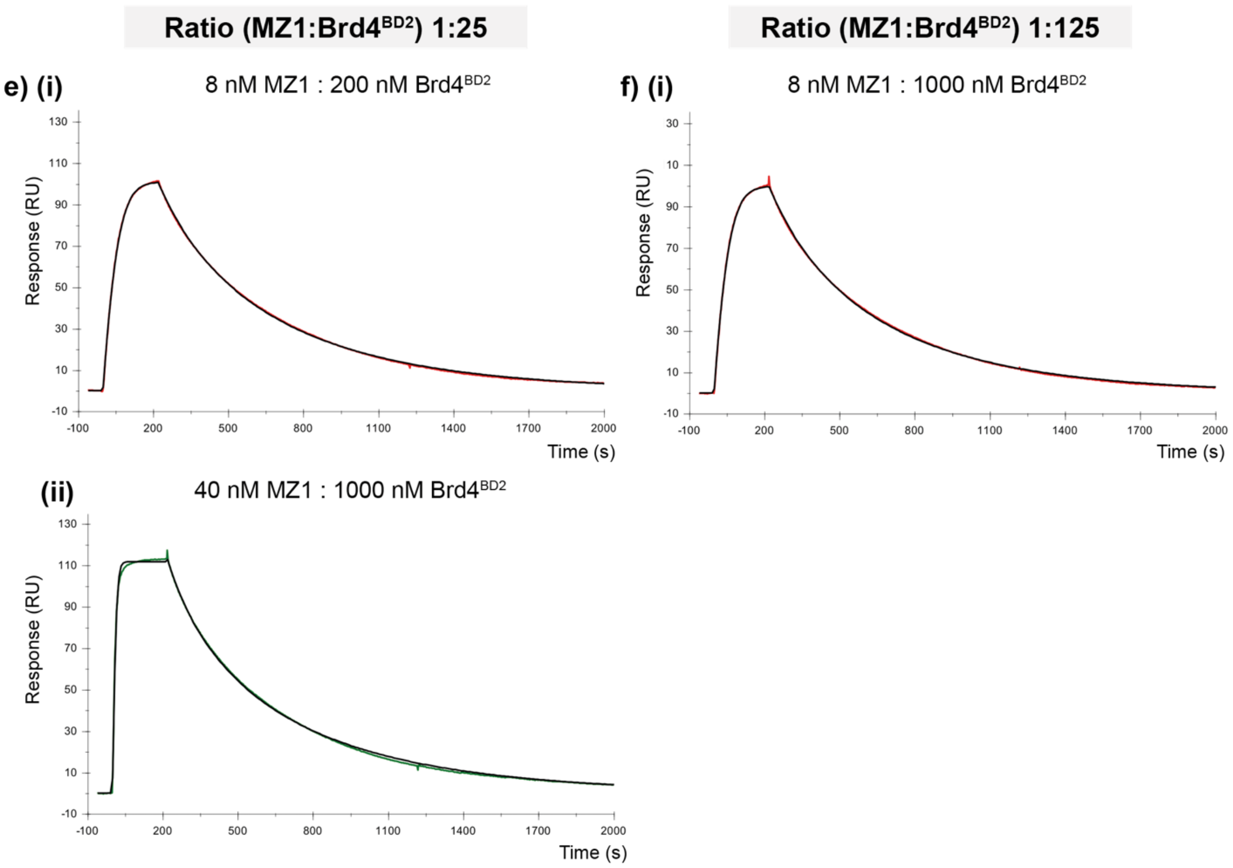


**Figure S5. Effect of varying the PROTAC:target ratio (MZ1:Brd4$^{BD2}$).** To further explore the anticipated effect of varying the ratio of the PROTAC and target protein on efficiency of ternary complex formation, a simple SPR study was undertaken whereby varying ratios of MZ1 and Brd4$^{BD2}$ (as depicted in (a)(i)), were mixed and allowed to equilibrate, then the binding response at 285.15 K measured by injecting over an SPR surface coated with immobilised biotin-VHL (~ 500 RU). For each injection, double-referenced sensorgrams were then fitted to a 1:1 Langmuir interaction model using Biacore T200 Evaluation Software (GE Healthcare). For wells in which the concentration of MZ1 would exceed that of Brd4$^{BD2}$ (dark grey squares depicted in (a)(i)), the binding data were not fitted, as in these cases the binding response for the MZ1:Brd4$^{BD2}$ complex (ternary complex formation with VHL) is reduced due to competitive binding of excess free PROTAC to form binary PROTAC:VHL complexes (the so-called 'hook effect'). For all other wells, binding constants are tabulated in (a)(ii). For different ratios of MZ1:Brd4$^{BD2}$ (1:1, 1:5, 1:25, 1:125), the fitted SPR binding constants ($K_D$, $k_{on}$, $k_{off}$, $R_{max}$) were then plotted (as shown in (b)). The fitted sensorgrams measured for each ratio (MZ1:Brd4$^{BD2}$) are shown in (c) - (f).

Although a limited study, the fitted binding data are consistent with our general expectations regarding selection of optimal PROTAC:target ratio for ternary binding studies (Figure S1). In particular, it is apparent that in this case use of a 1:1 ratio of PROTAC:target would seem likely to result in underestimation of the true binding affinity ($K_D$) for the ternary complex (refer (a)(i)). The fitted on-rate is slowest for the 1:1 ratio (MZ1:Brd4$^{BD2}$) (plot of $k_{on}$, in (a)(ii)) and the fitted off-rate also is fastest for this ratio (plot of $k_{off}$, in (a)(i)). Similarly, the fitted $R_{max}$ appears to be lower for the 1:1 ratio (refer (b)(iv)). This is consistent with our other kinetic data (Table S2) indicating that MZ1 both binds more slowly to VHL and dissociates more quickly, as compared to the interaction of the MZ1:Brd4$^{BD2}$ complex with VHL. As the measured SPR binding response of a mixture of MZ1 and MZ1:Brd4$^{BD2}$ will be dominated by the significantly higher molecular weight of MZ1:Brd4$^{BD2}$ (relative to MZ1 alone), the most apparent effects of free MZ1 competing for binding might be expected to be a reduction in the overall binding response and a shift in the fitted kinetic parameters for MZ1:Brd4$^{BD2}$ binding towards those of MZ1. These data are consistent with this conclusion.

Notably, as the ratio of Brd4$^{BD2}$ relative to MZ1 increases, each of the fitted kinetic parameters appear to gradually coalesce (refer plots in (b)). Again, this is likely due the reduced concentrations of free MZ1 remaining in the injected well solution, such that its competitive effect on the measured binding response is gradually reduced, until essentially only ternary complex formation is measured.

**Binding to immobilised Brd4$^{BD2}$ (~400 RU)**

**a) MZ1**

**Binary**

| $k_{on}$ (1/Ms) (x $10^5$) | $k_{off}$ (1/s) | $K_D$ (nM) | $R_{max}$ (RU) |
|---|---|---|---|
| 26 | 0.020 | 7.5 | 17.2 |

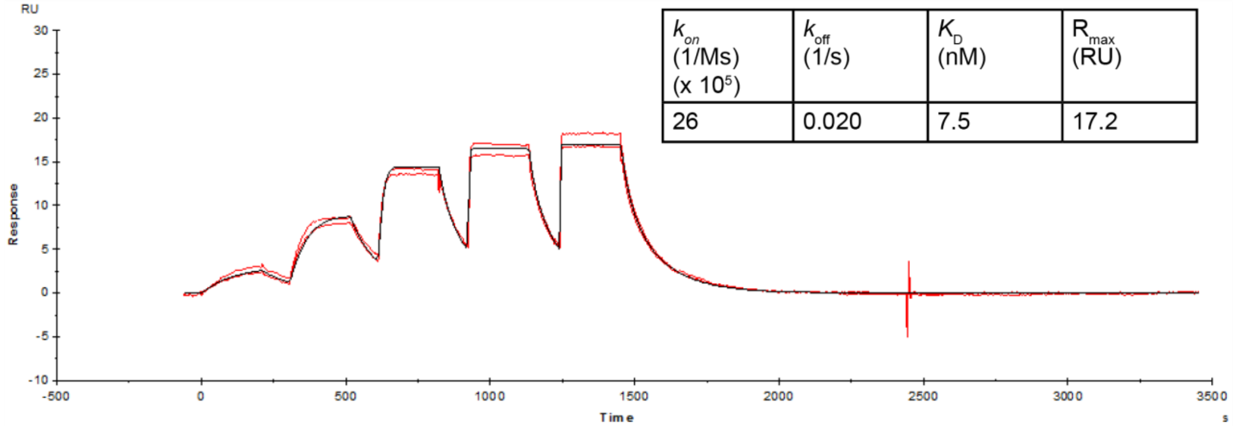


**b) MZ1:VHL**

**Ternary**

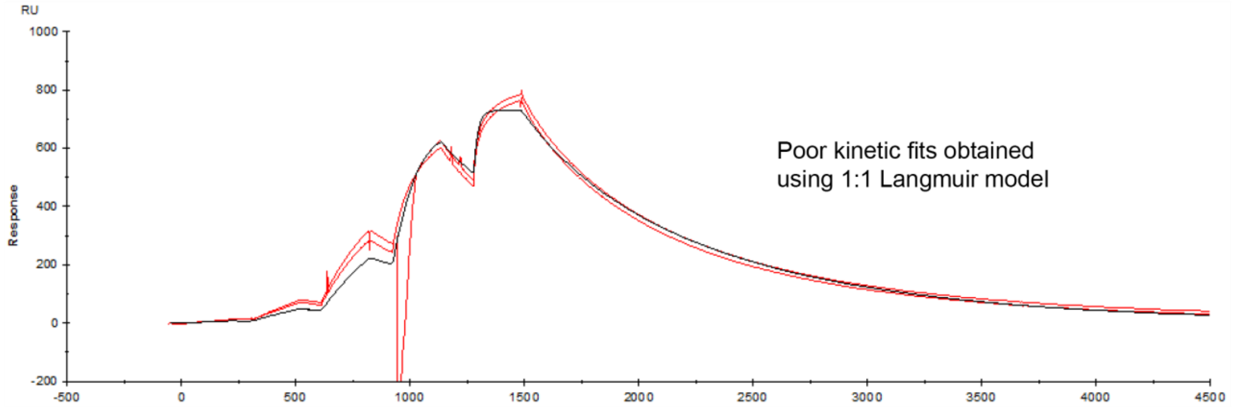


[Figure S6 continues on the next page.]

### Binding to immobilised Brd4$^{BD2}$ (~400 RU)

**c) AT1** **Binary**

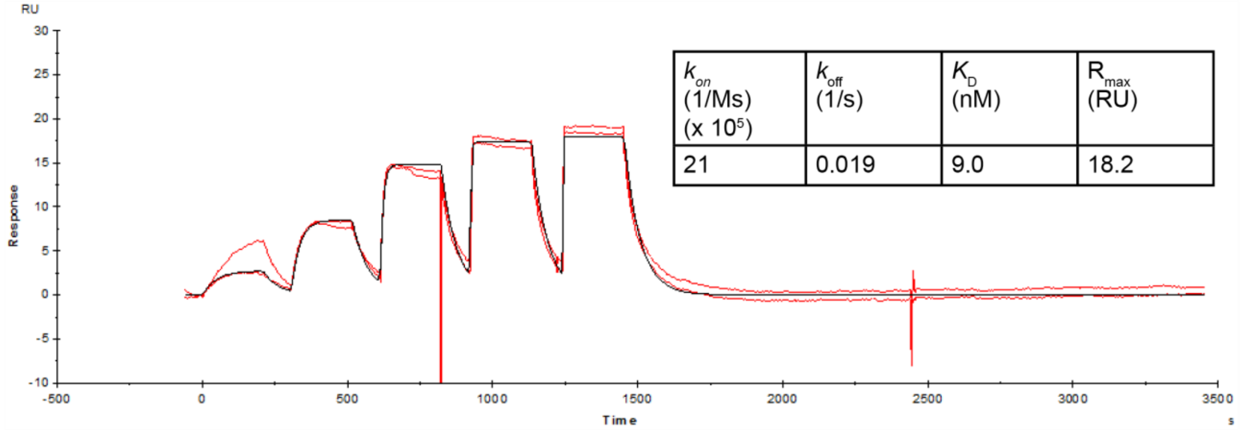


| $k_{on}$ (1/Ms) (x $10^5$) | $k_{off}$ (1/s) | $K_D$ (nM) | $R_{max}$ (RU) |
|---|---|---|---|
| 21 | 0.019 | 9.0 | 18.2 |

**d) AT1:VHL** **Ternary**

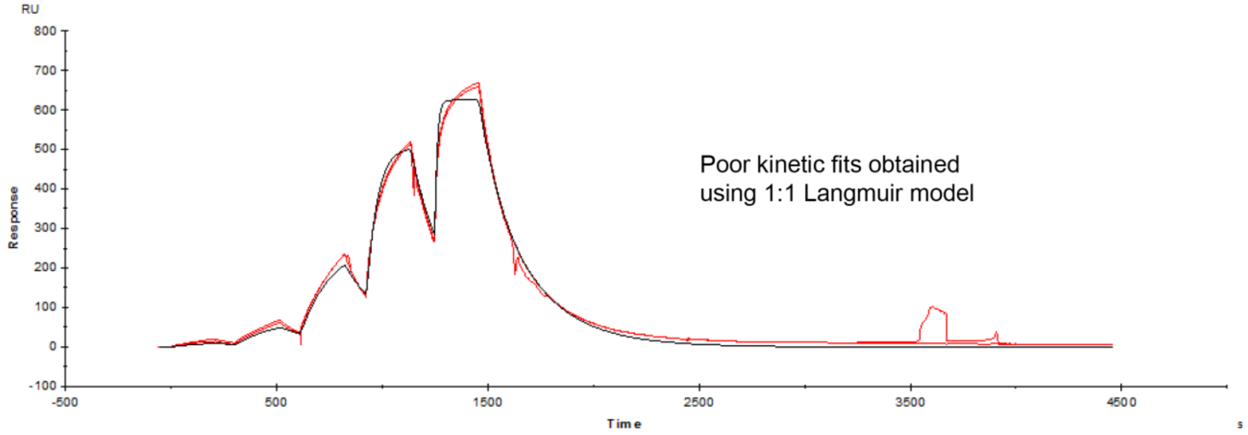


[Figure S6 continues on the next page.]

### Binding to immobilised Brd4$^{BD2}$ (~400 RU)

### e) MZP61      <u>Binary</u>

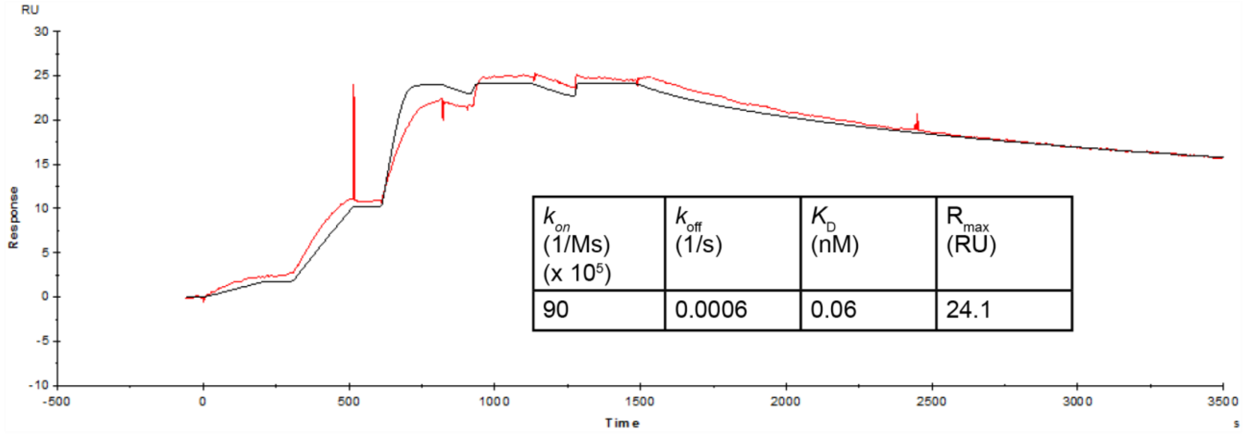


| $k_{on}$ (1/Ms) (x $10^5$) | $k_{off}$ (1/s) | $K_D$ (nM) | $R_{max}$ (RU) |
|---|---|---|---|
| 90 | 0.0006 | 0.06 | 24.1 |

### f) MZP61:VHL      <u>Ternary</u>

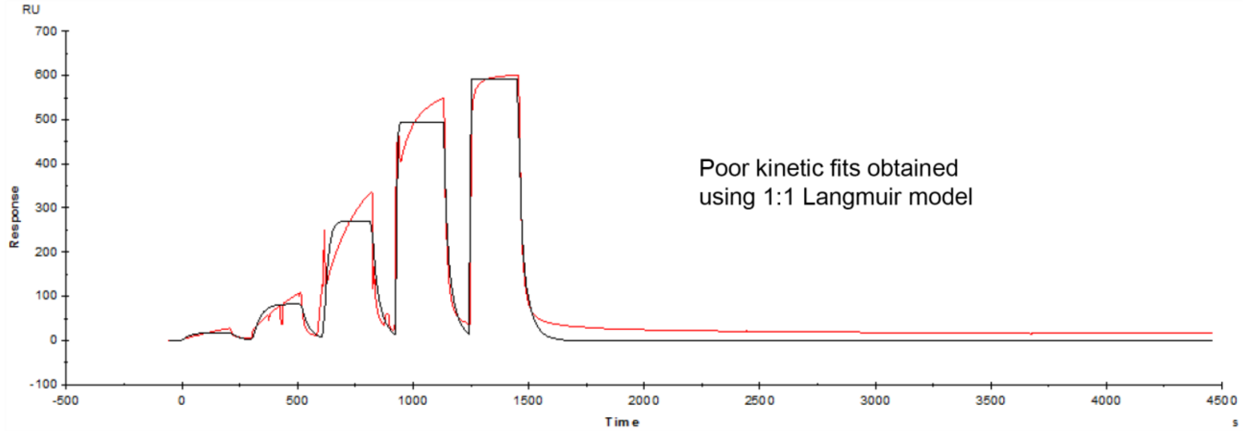


**Figure S6. Reversed-format SPR binding experiments (immobilised Brd4$^{BD2}$).** SPR sensorgrams are shown for preliminary experiments conducted in a reversed format, for PROTAC (binary) or PROTAC:VHL (ternary) binding to immobilised Brd4$^{BD2}$ (for PROTACs MZ1, AT1, MZP61), measured at 285.15 K. These data are shown for qualitative, rather than quantitative purposes. Whilst reasonable kinetic fits to a 1:1 Langmuir interaction model were obtained for binary binding data (in particular for MZ1 and AT1), this was not the case for ternary experiments. Although all ternary binding experiments show a significant increase in overall binding response consistent with binding of the anticipated PROTAC:VHL complex (relative to PROTAC alone), the resulting ternary sensorgrams were not able to be adequately fitted using a 1:1 Langmuir interaction model. Qualitatively, however, a number of observations are able to be made. Firstly, despite the relatively poor kinetic fitting, it is notable that the binary binding of MZP61 to immobilised Brd4$^{BD2}$ appears to be high-affinity, characterised by very slow dissociation kinetics (refer (e)) as compared to either MZ1 (refer (a)) or AT1 (refer (c)). This is indeed consistent with the higher expected binary affinity of the tetrahydroisoquinoline binder of MZP61 for Brd4$^{BD2}$, relative to the triazolodiazepine binder of either MZ1 or AT1. [2] Secondly, in the presence of VHL, the rate of dissociation of the MZP61:VHL complex from immobilised Brd4$^{BD2}$ (refer (f)) can be observed to be qualitatively very much faster than MZP61 alone, reflecting the overall negative cooperativity expected for this ternary complex relative to the binary interaction.

### Binding to immobilised biotin-VHL:

#### Ternary

#### a) MZ1:Brd2$^{BD1}$

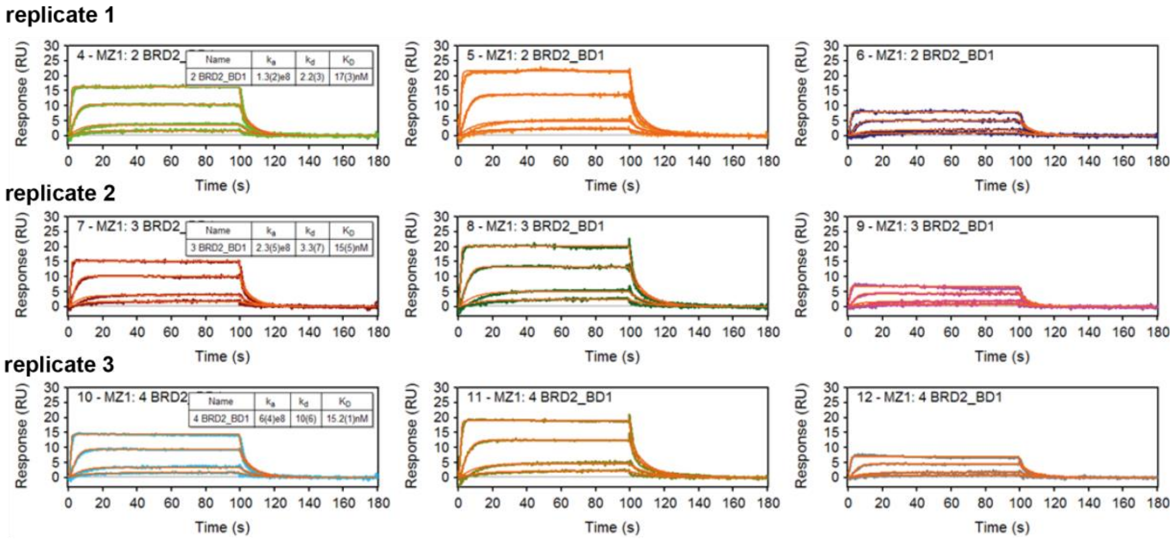


#### b) MZ1:Brd2$^{BD2}$

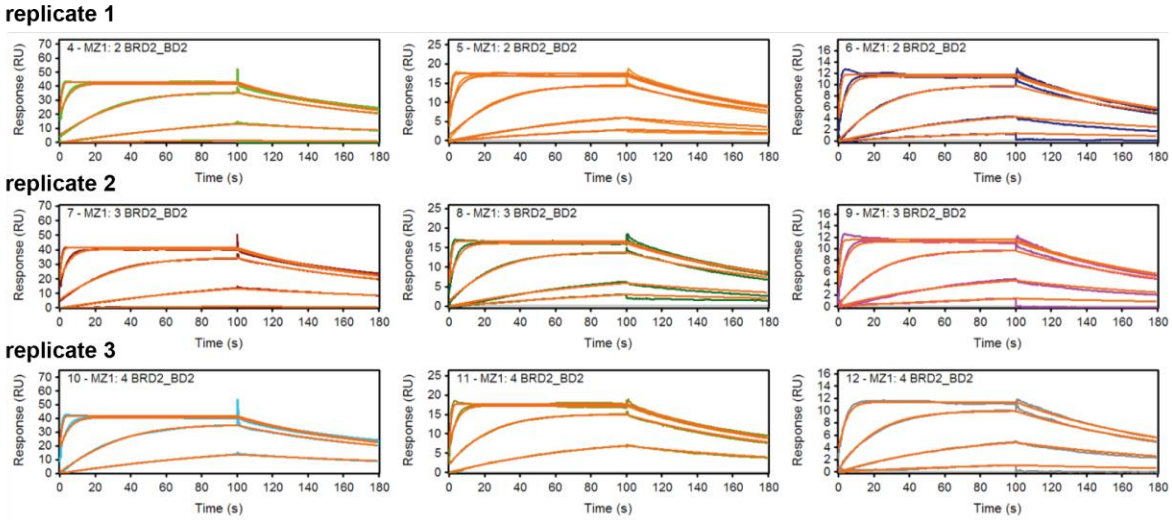


[Figure S7 continues on the next page.]

**Binding to immobilised biotin-VHL:**

**<u>Ternary</u>**

### c) MZ1:Brd3$^{BD1}$

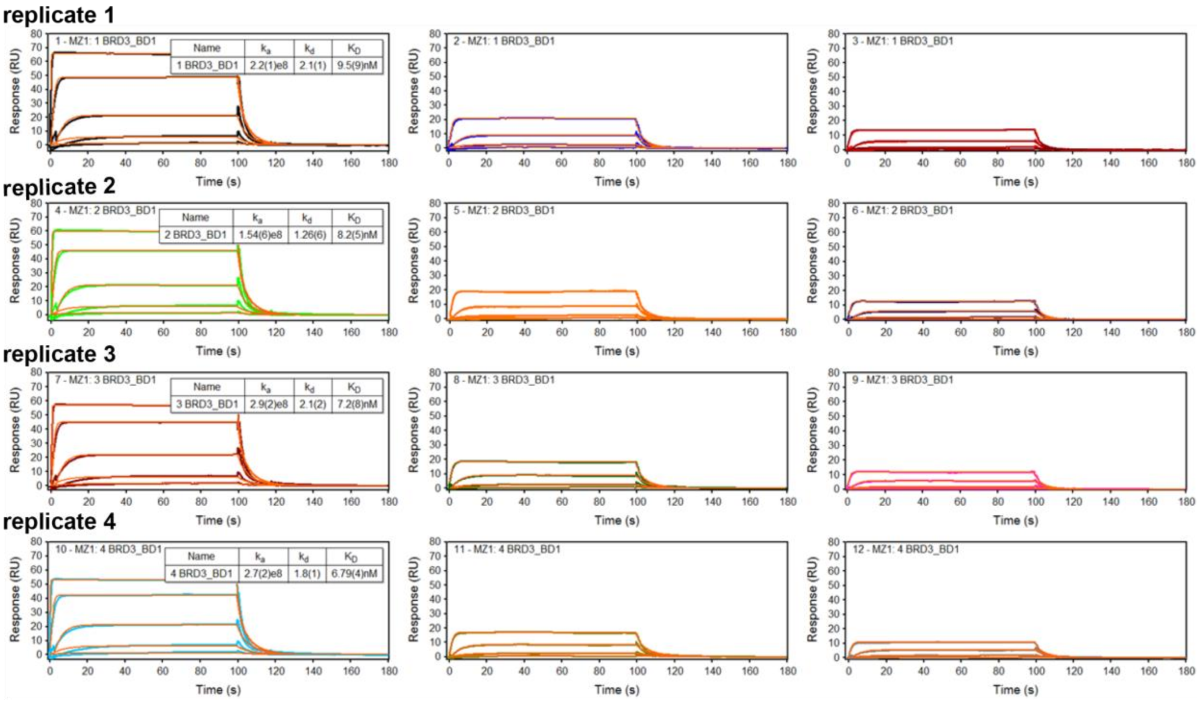


### d) MZ1:Brd3$^{BD2}$

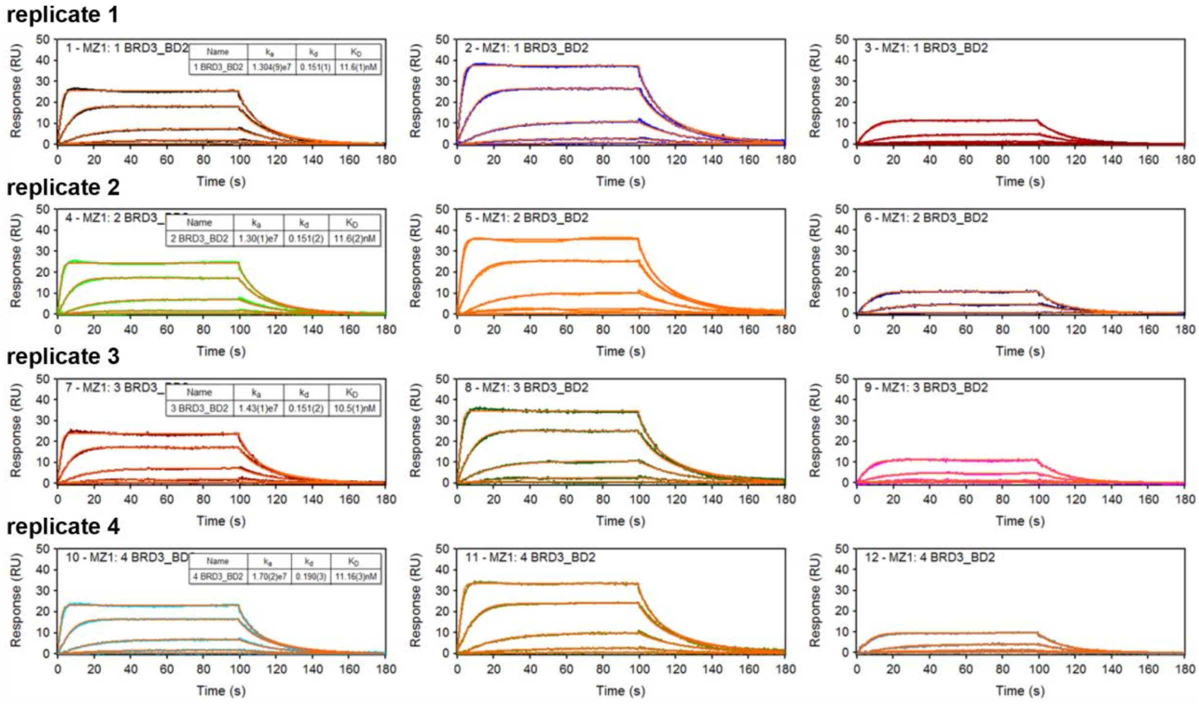


[Figure S7 continues on the next page.]

### Binding to immobilised biotin-VHL:

**<u>Ternary</u>**

#### e) MZ1:Brd4$^{BD1}$

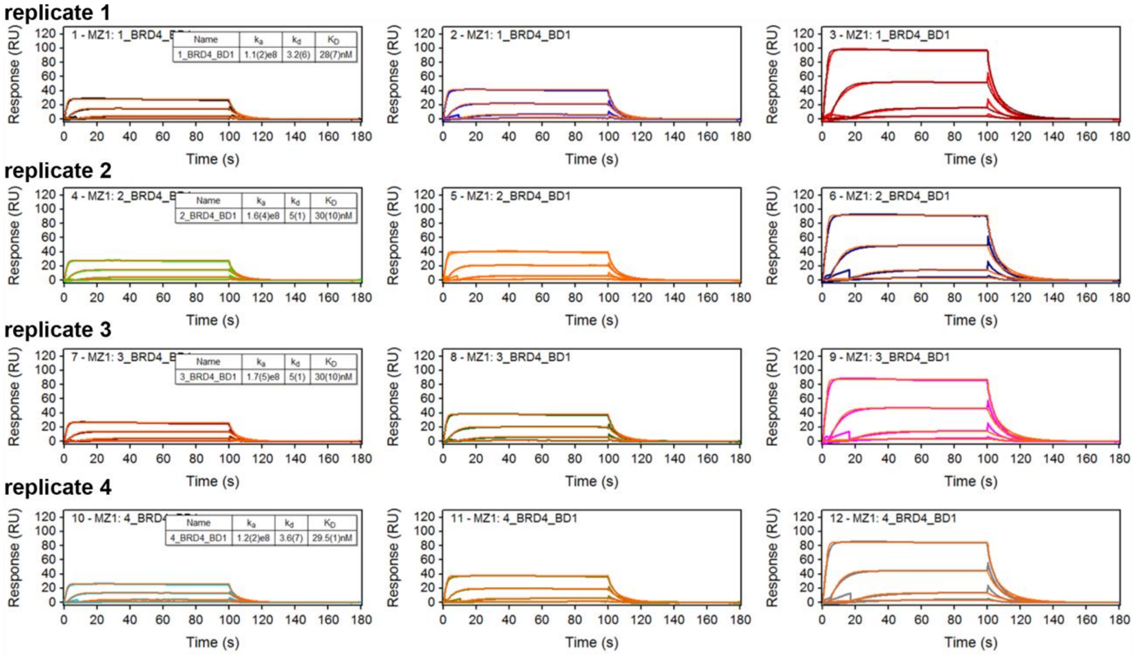


#### f) MZ1:Brd4$^{BD2}$

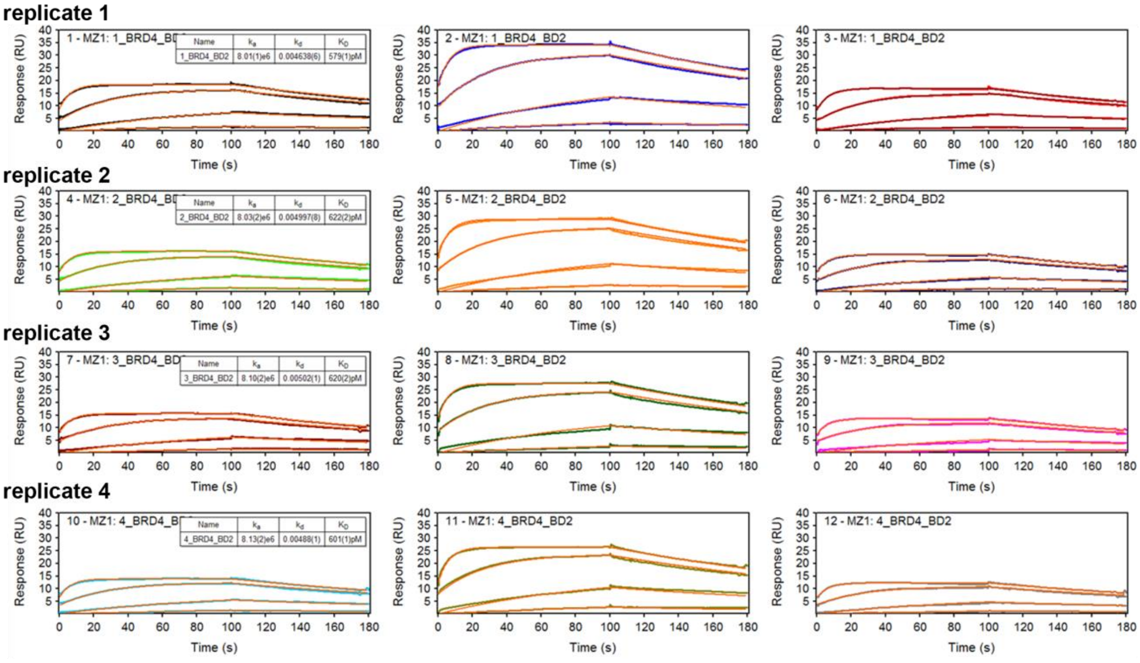


**Figure S7. Representative SPR sensorgrams for MZ1:BD (ternary) binding to immobilised VHL (varying the individual BET bromodomain).** SPR binding analyses for ternary complexes were performed in single-cycle kinetic (SCK) mode at 298.15 K and analysed as described (Figure S3).

**Binding to immobilised biotin-VHL:**

**<u>Ternary</u>**

### a) AT1:Brd2$^{BD1}$

**replicate 1**

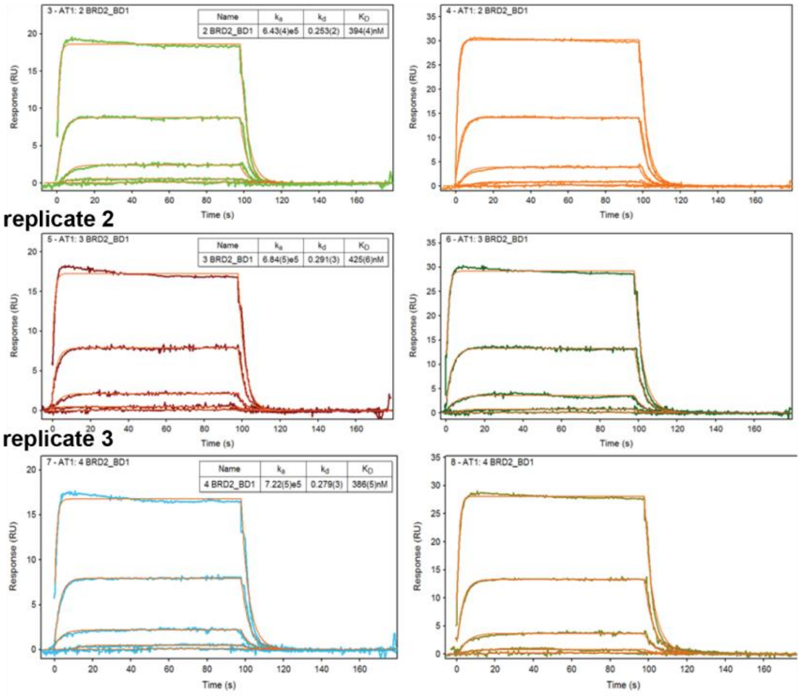


### b) AT1:Brd2$^{BD2}$

**replicate 1**

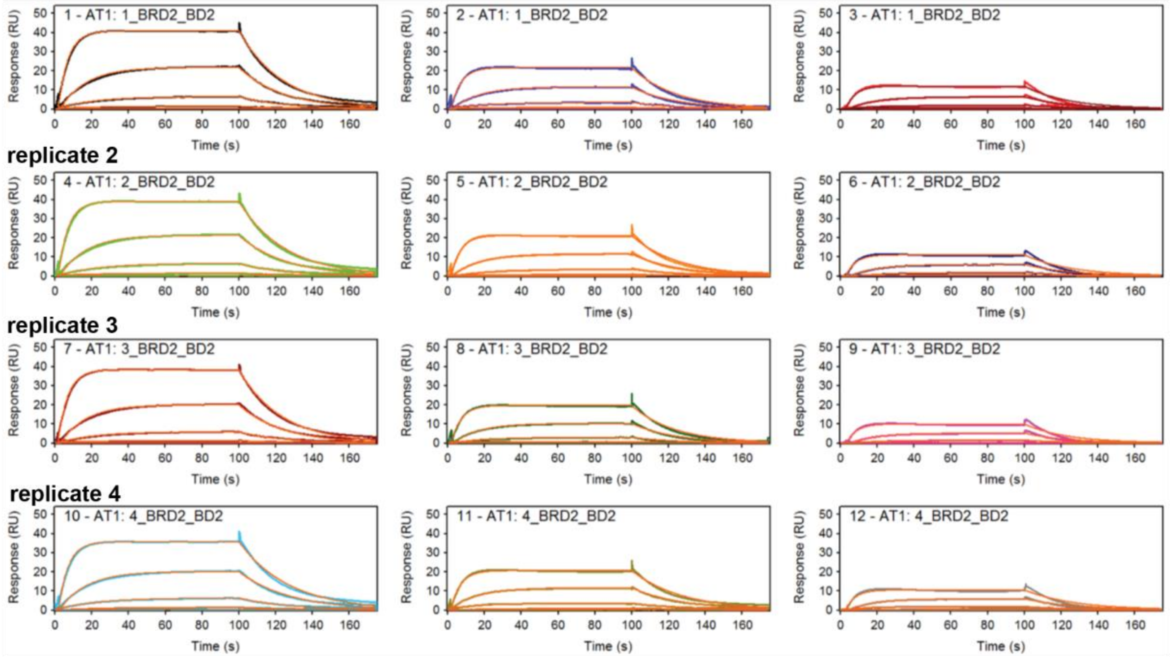


[Figure S8 continues on the next page.]

**Binding to immobilised biotin-VHL:**

**<u>Ternary</u>**

**c) AT1:Brd3$^{BD1}$**

**replicate 1**

**replicate 2**

**replicate 3**

**replicate 4**

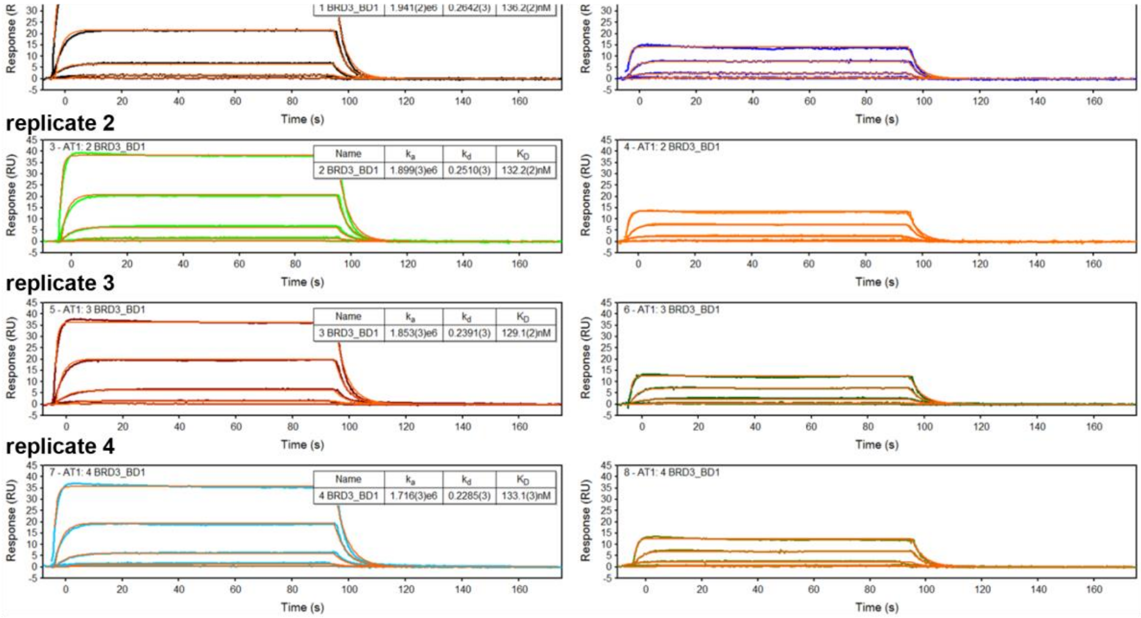


**d) AT1:Brd3$^{BD2}$**

**replicate 1**

**replicate 2**

**replicate 3**

**replicate 4**

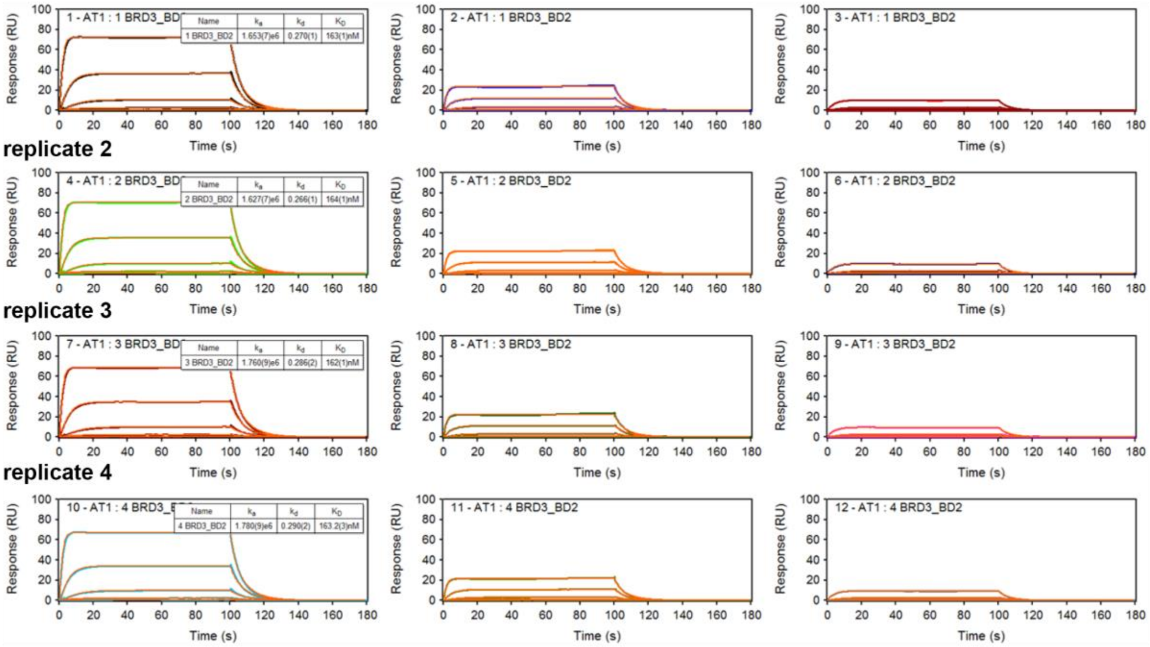


[Figure S8 continues on the next page.]

**Binding to immobilised biotin-VHL:**

**Ternary**

### e) AT1:Brd4$^{BD1}$

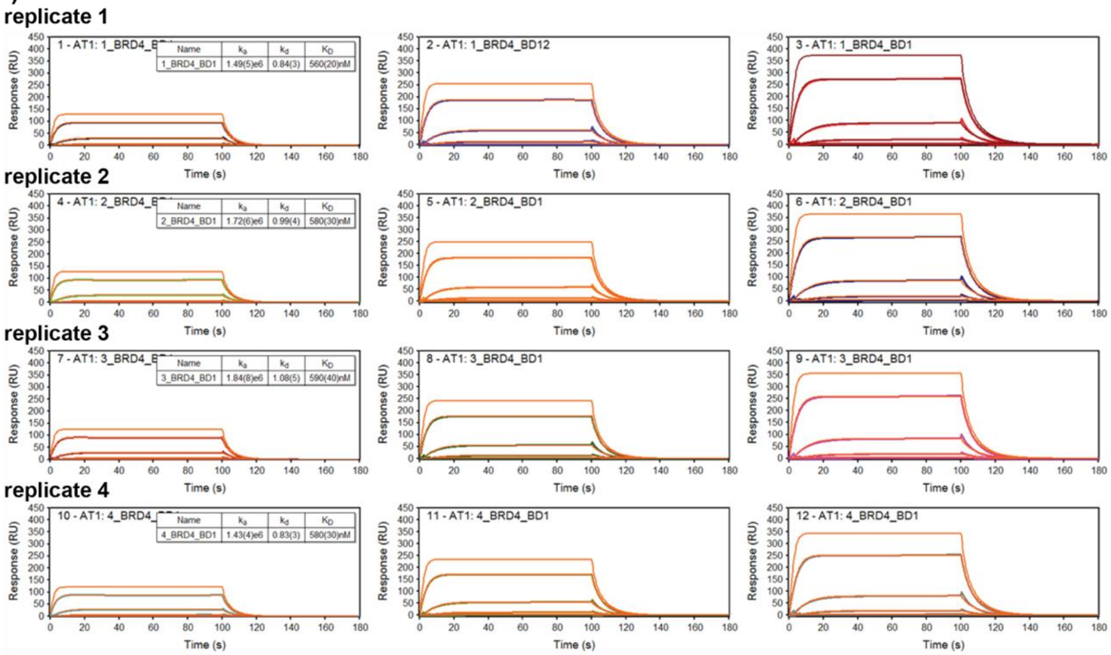


### f) AT1:Brd4$^{BD2}$

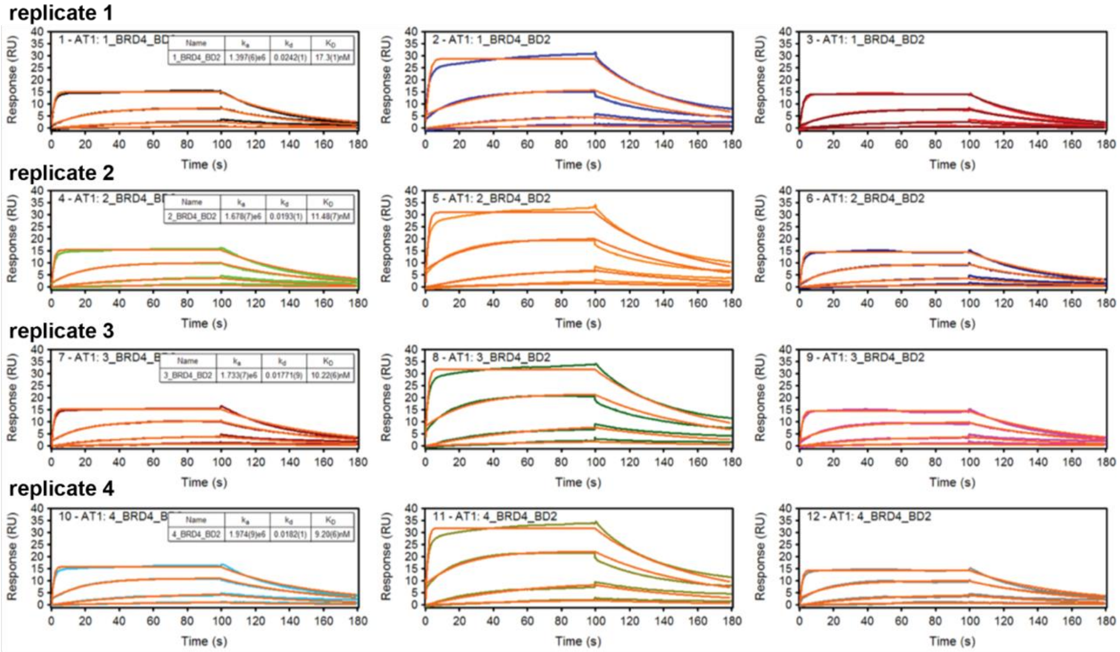


**Figure S8. Representative SPR sensorgrams for AT1:BD (ternary) binding to immobilised VHL (varying the individual BET bromodomain).** SPR binding analyses for ternary complexes were performed in single-cycle kinetic (SCK) mode at 298.15 K and analysed as described (Figure S3).

**Binding to immobilised biotin-VHL:**

**<u>Ternary</u>**

#### a) MZ1:Brd2$^{BD2,G382E}$

**replicate 1**

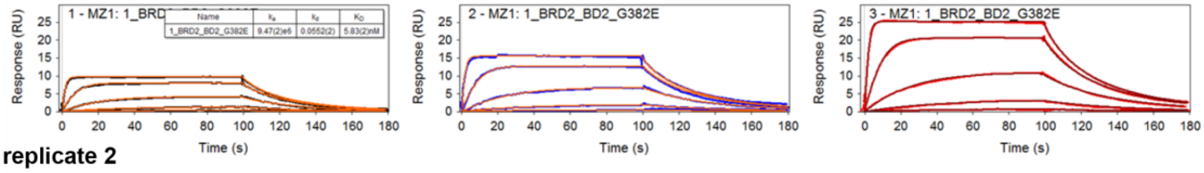


**replicate 2**

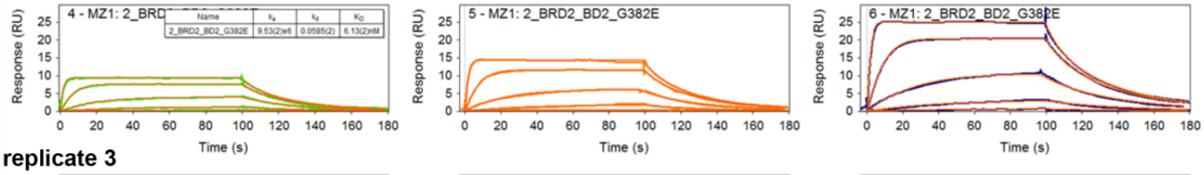


**replicate 3**

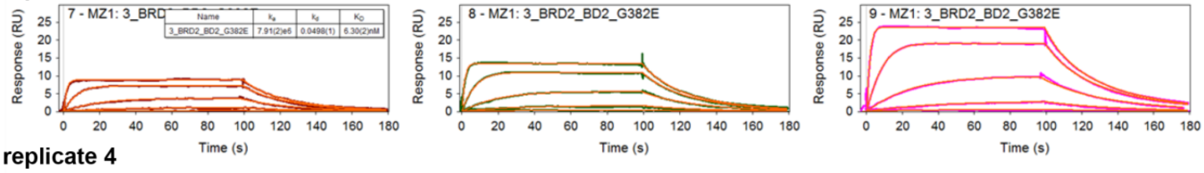


**replicate 4**

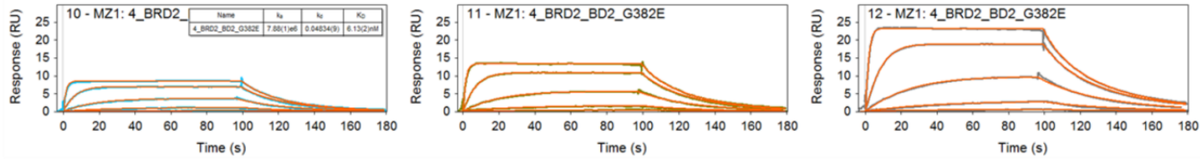


#### b) AT1:Brd2$^{BD2,G382E}$

**replicate 1**

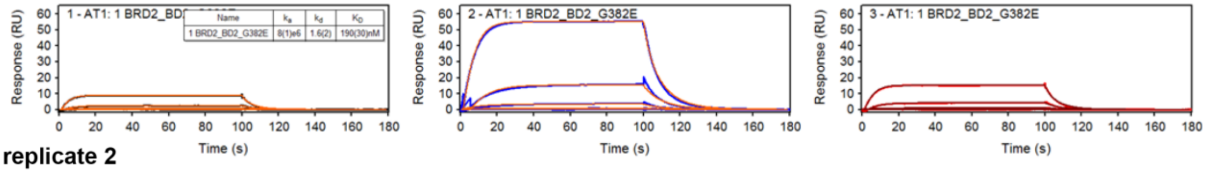


**replicate 2**

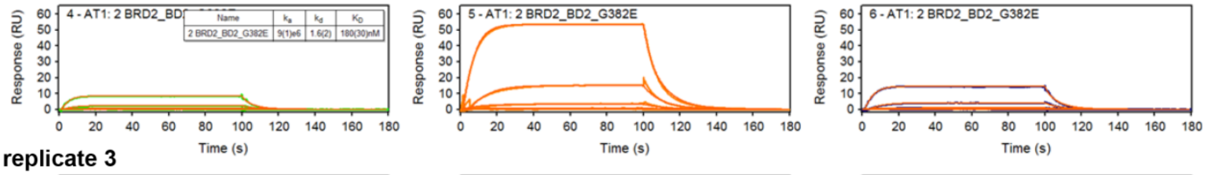


**replicate 3**

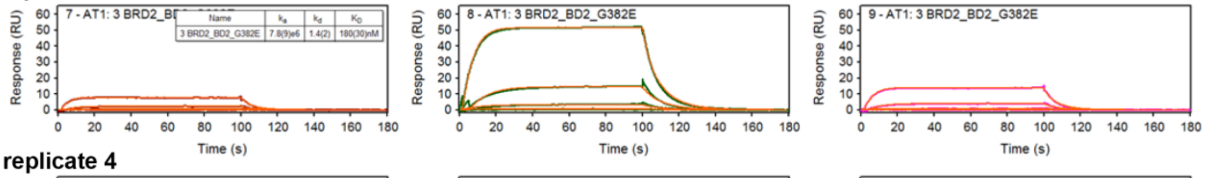


**replicate 4**

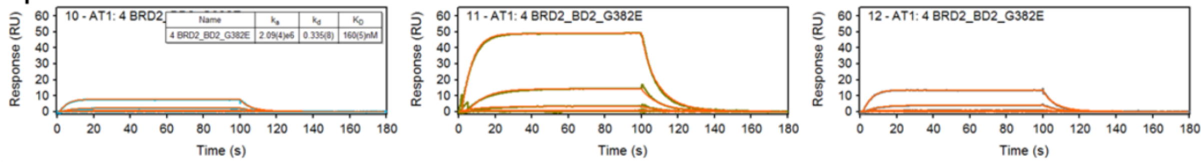


[Figure S9 continues on the next page.]

**Binding to immobilised biotin-VHL:**

**<u>Ternary</u>**

#### c) MZ1:Brd3$^{BD2,E344G}$

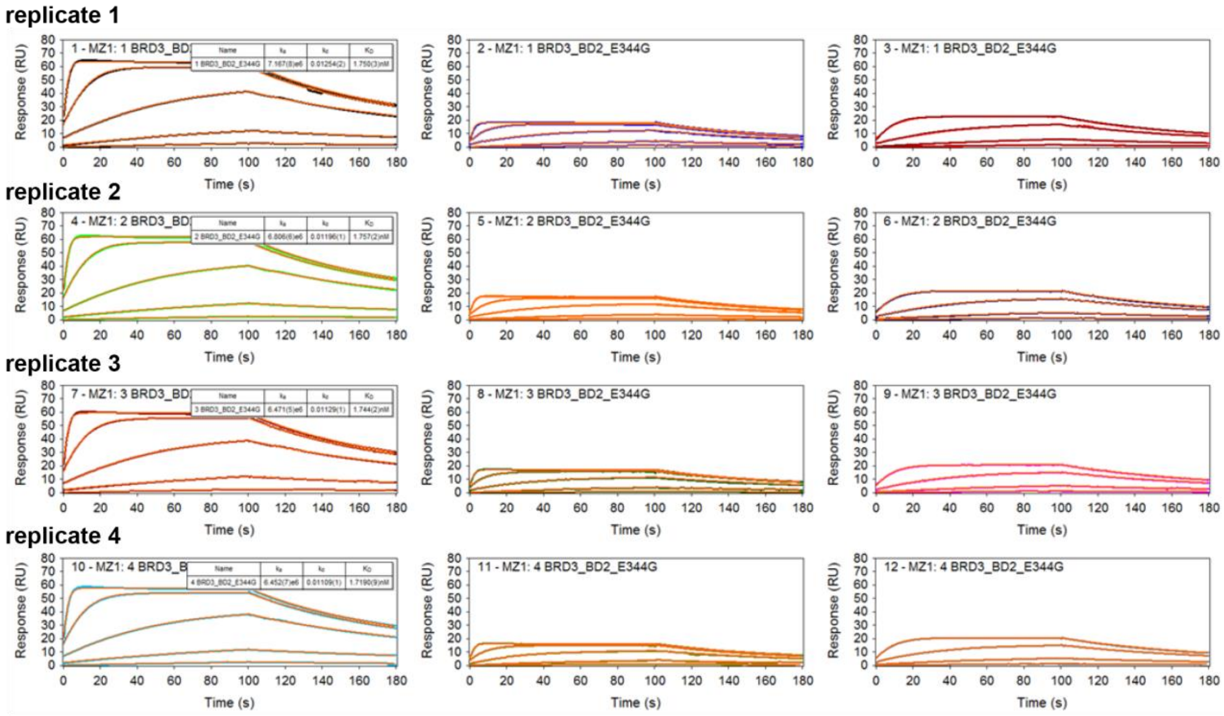


#### d) AT1:Brd3$^{BD2,E344G}$

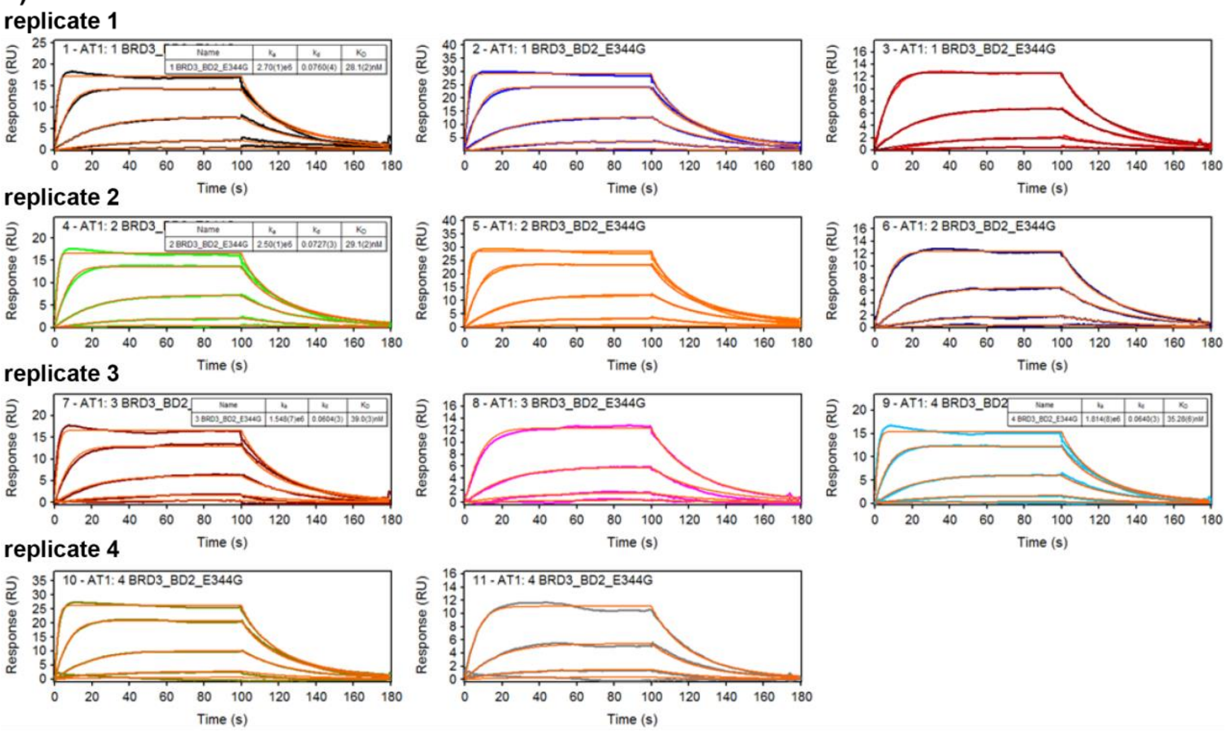


[Figure S9 continues on the next page.]

### Binding to immobilised biotin-VHL:

**Ternary**

#### e) MZ1:Brd4$^{BD2,G386E}$

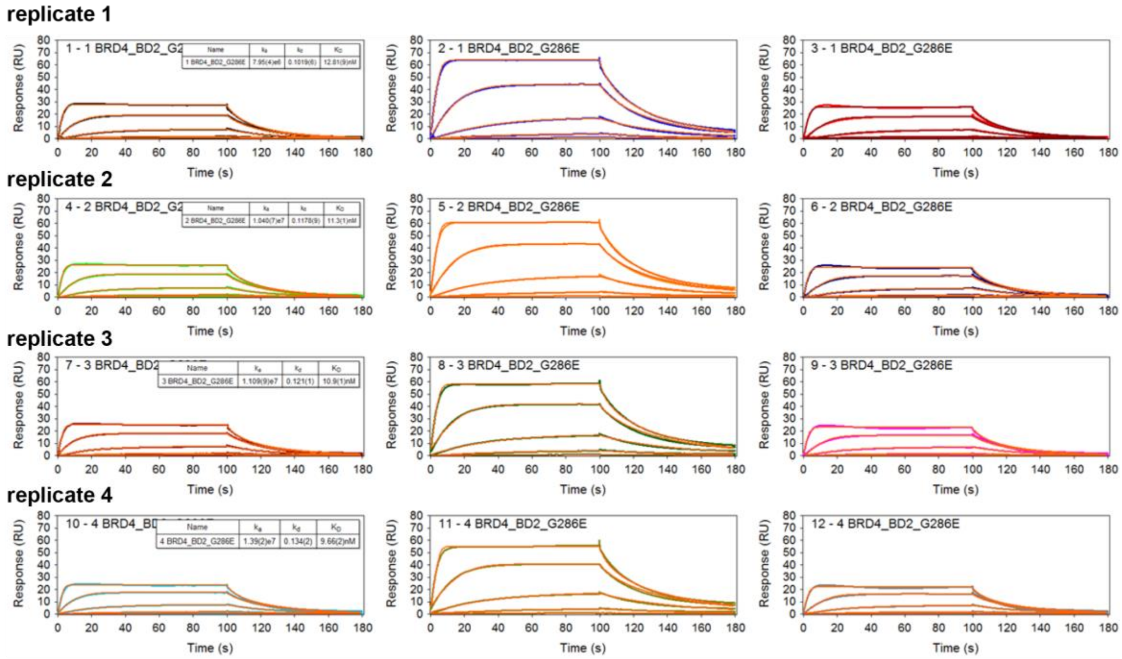


#### f) AT1:Brd4$^{BD2,G386E}$

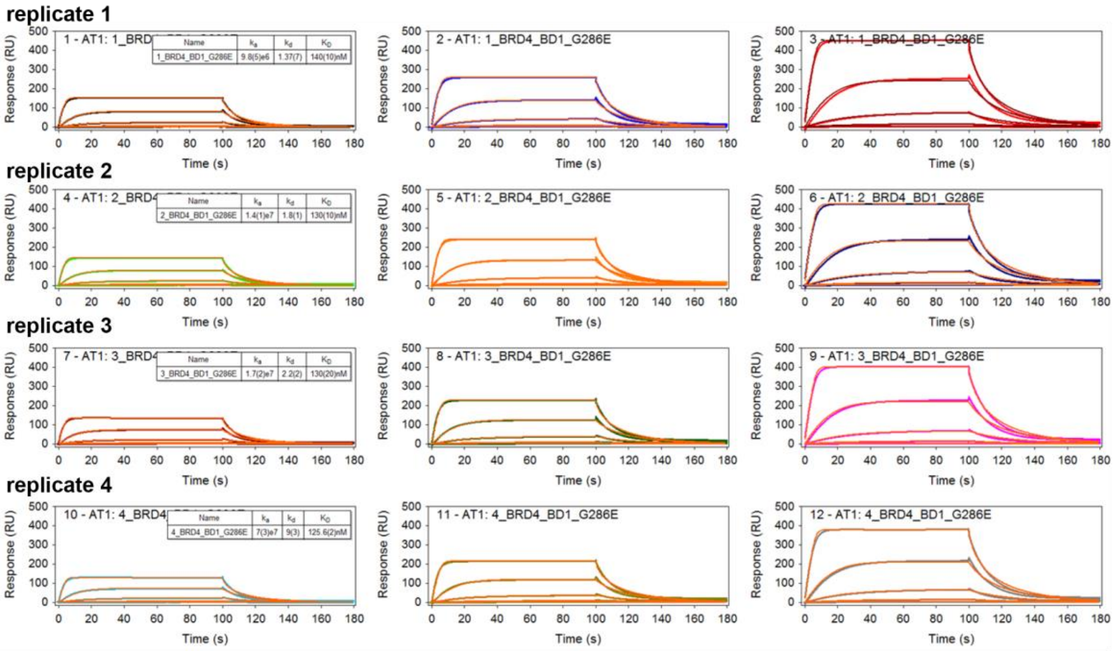


**Figure S9. Representative SPR sensorgrams for PROTAC:BD (ternary) binding to immobilised VHL (for PROTACs MZ1 and AT1, and different BET bromodomain point-mutants).** SPR binding analyses for ternary complexes were performed in single-cycle kinetic (SCK) mode at 298.15 K and analysed as described (Figure S3).

**a) MZ1 (binary) or MZ1:BD (ternary) for individual BET bromodomains.**

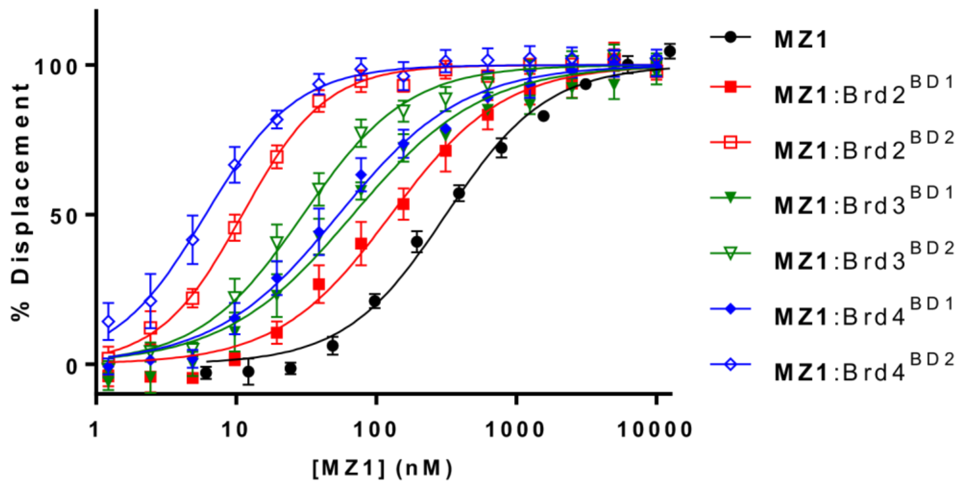


**b) MZ1 (binary) or MZ1:BD (ternary) for BET bromodomain point-mutants**

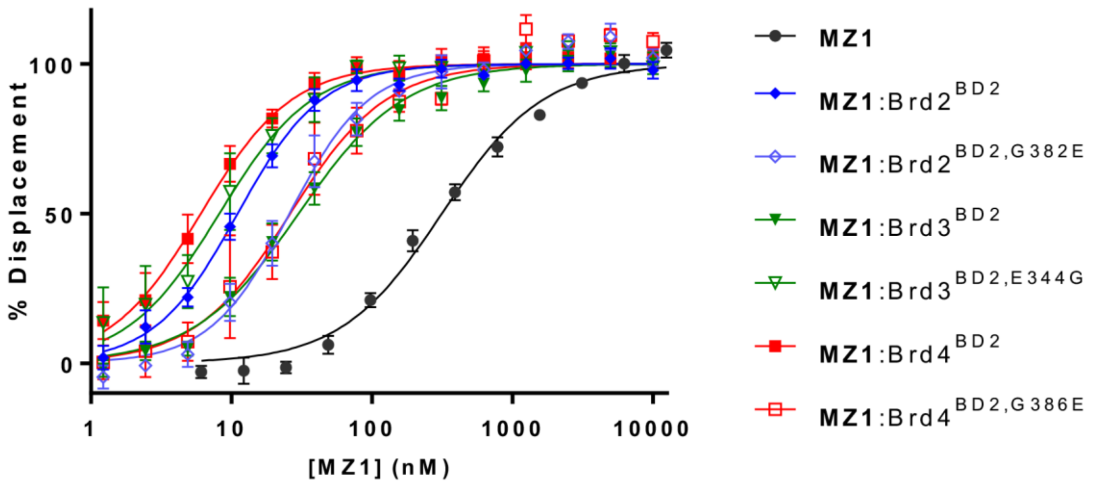


**Figure S10. Fitted Fluorescence Polarization (FP) competition data for MZ1 (binary) and MZ1:BD (ternary) binding to VHL in solution (for individual BET bromodomains and bromodomain point-mutants).**

Plotted data represents mean percent displacement of fluorescent HIF-1$\alpha$ probe molecule from VHL (MZ1 alone or MZ1 + saturating concentration of BD), from three independent experiments (errors are SD, N=3 for A, N=2 for B), fitted using non-linear regression to determine half-maximal inhibitory concentration ($IC_{50}$) values.

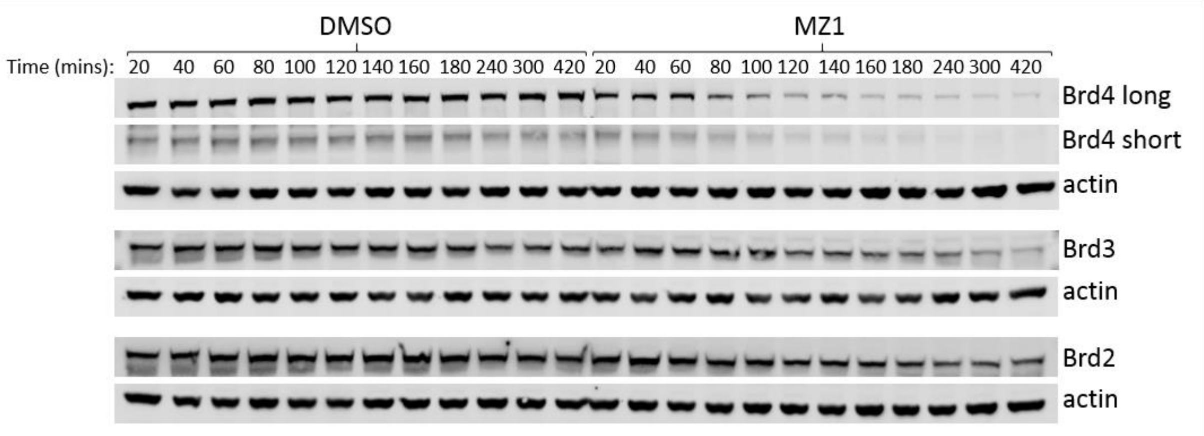


**Figure S11. Representative Western blot for degradation time course data using HEK293 cells.**

Cells were treated with 0.1% v/v DMSO (vehicle) or 333 nM MZ1 over a range of time points (20 min – 420 min) prior to lysis. Samples (40 μg total protein/well) were resolved by SDS-PAGE, transferred to nitrocellulose membrane and probed with anti-Brd2, anti-Brd3 or anti-Brd4 primary antibodies, followed by either goat anti-mouse or donkey anti-rabbit IRDye 800CW secondary antibodies. Bands were then detected using a ChemiDoc (BioRad) and quantified (Image Studio Lite, version 5.2) with normalisation to β-actin and DMSO control per time point.

**Supporting Methods**

### 1. Constructs, protein expression and purification

Wild-type and mutant versions of human proteins were used for all protein expression, as follows: VHL (UniProt accession number: P40337), ElonginC (Q15369), ElonginB (Q15370) and the bromodomains (BDs) of Brd2 (P25440), Brd3 (Q15059), and Brd4 (O60885). Synthetic DNA gene fragments and oligonucleotide primers were purchased from were purchased from Integrated DNA Technologies (IDT) and Sigma Aldrich respectively.

#### BET bromodomains (BDs), point-mutants and biotinylated Brd4$^{BD2}$

Plasmids (pNIC28-Bsa4 Kan$^{r}$) containing the single BET bromodomain constructs - Brd2$^{BD1}$ (71-194), Brd2$^{BD2}$ (344-455), Brd3$^{BD1}$ (24-144), Brd3$^{BD2}$ (306-416), Brd4$^{BD1}$ (44-168), and Brd4$^{BD2}$ (333-460) in frame with an N-terminal His$_6$-tag and TEV protease cleavage site were provided by the Oxford Structural Genomics Consortium (SGC).[1, 4] Single-point bromodomain mutations were introduced using *QuikChange II Site directed Mutagenesis Kit* (Agilent), as described in detail previously,[4] with the modification that digestion of parental DNA was achieved using FastDigest DpnI (Thermo Fisher) for 1 h at 37 °C. DNA from single-colony clones was extracted and purified using the Monarch Plasmid Miniprep Kit (NEB) and submitted for sequencing (MRC PPU Reagents and Services facility, University of Dundee, Scotland) to confirm the presence of the desired mutation.

Individual BET bromodomains and bromodomain point mutants were expressed and purified as described previously,[1, 4] with the following modifications. The clarified cell lysate was affinity purified using His Trap HP (1mL) Ni sepharose columns (GE Healthcare) and eluted in 250 mM imidazole in 20 mM 4-(2-hydroxyethyl)-1-piperazineethanesulfonic acid (HEPES), 500 mM sodium chloride and 1 mM *β*-mercaptoethanol, pH 7.5. Eluted proteins were purified directly without cleavage of the His$_6$ tag, by size exclusion chromatography (SEC) on a Superdex 75 16/60 Hiload gel filtration column on an ÄKTApure™ system (GE Healthcare) in the following buffer: 20 mM HEPES, 500 mM sodium chloride and 1 mM tris(2-carboxyethyl)phosphine (TCEP), pH 7.5. The mass and purity of the proteins were subsequently verified by mass spectrometry (FingerPrints Proteomics Facility, University of Dundee, Scotland).

*Chemical biotinylation of Brd4$^{BD2}$.*

To chemically biotinylate Brd4$^{BD2}$, the protein was mixed in a 1:1 stoichiometric ratio with EZ-Link NHS-PEG4-biotin (Thermo Scientific) and incubated at room temperature for 1 h. Unreacted NHS-biotin was removed by passing the sample over a PD Minitrap G-25 desalting column (GE Healthcare) into 20 mM HEPES, 500 mM sodium chloride and 1 mM tris(2-carboxyethyl)phosphine (TCEP), pH 7.5.

**VHL and biotinylated VHL.**

*Cloning of VCB-AviTag complex and expression and purification of VCB and VCB-AviTag complexes.*

A synthetic DNA sequence (gBlock) was purchased from IDT, which encoded for ElonginB (1-104) followed by a short spacer and AviTag sequence at the C-terminus (final translated protein sequence: MDVFLMIRRHKTTIFTDAKESSTVFELKRIVEGILKRPPDEQRLYKDDQLLDDGKTLGECGFTSQTARPQAPATVGLAFR ADDTFEALCIEPFSSPPELPDVMKgspagggIndifeaqkiewhe). This DNA was sub-cloned into the NcoI/HindIII sites of a plasmid (pIVM02, pCDFDUET-1b, Strep$^r$),[5] which already contained a sequence that encoded for ElonginC (17-112). The final plasmid was co-transformed into BL21(DE3) *E. coli* cells along with the plasmid for expression of VHL (54-213) with an N-terminal His6 purification tag and TEV cleavage site (pHAT4,[6] Amp$^r$). Both VCB (VHL$^{54-213}$:ElonginC$^{17-112}$:ElonginB$^{1-104}$) and the VCB-AviTag complexes (VHL$^{54-213}$:ElonginC$^{17-112}$:ElonginB$^{1-104}$-AviTag) were co-expressed and purified, including removal of the His6 tag using TEV protease, as previously described for the VCB complex.[1] Both purified complexes were stored in 20 mM HEPES, 100 mM sodium chloride and 1 mM TCEP, pH 7.5.

*Expression and purification of GST-BirA.*

A plasmid (pGEX6P-1, Amp$^r$) containing the BirA enzyme as an N-terminal GST-fusion protein with a TEV protease cleavage site (gift of the MRC PPU Reagents and Services, University of Dundee, Scotland; Genbank: M10123) was transformed into BL21 (DE3) cells and expressed and purified based on a modified literature procedure.[7] Briefly, a 10 mL starter culture of LB medium containing ampicillin (100 µg/mL) and D-glucose (0.4% v/v) was inoculated from a single colony and grown overnight at 37 °C in a shaking incubator (200 rpm). The starter culture (8 mL) was added a 1 L culture of TB containing ampicillin (100 µg/mL) and D-glucose (0.8 % v/v) and grown at 37 °C for 3 h. At an optical density ($A_{600}$) of approximately 1.1, the temperature was lowered to 23 °C and expression was induced using Isopropyl β-D-1-thiogalactopyranoside (IPTG) (0.4 mM) for approximately 16 h at 23 °C (180 rpm). Cells were harvested by centrifugation (20 min, 4200 rpm) in a JC-M6 centrifuge (Beckman Coulter). Cells were resuspended on ice in 50 mL of GST buffer consisting of 50 mM HEPES, 500 mM sodium chloride, 5% v/v glycerol and 5 mM DTT, supplemented with Complete protease inhibitor (Roche) and lysed using a Stansted Cell Disruptor (Stansted Fluid Power). Lysate was centrifuged (20,000 rpm, 4 °C) in an Avanti J-25 centrifuge (Beckman Coulter), filtered (0.45 µM syringe filter) and passed twice over a Glutathione Sepharose 4B resin (5 mL bed volume) (GE Healthcare) pre-equilibrated in GST Buffer. The column was washed with GST Buffer (40 mL) and the protein eluted in GST Buffer containing 20 mM L-glutathione (Sigma Aldrich) (20 mL). The eluted GST-BirA protein was purified directly by SEC on a Superdex 75 16/60 Hiload gel filtration column on an ÄKTApureTM system (GE Healthcare) in the following buffer: 20 mM HEPES, 150 mM sodium chloride, pH 7.5 and the protein concentrated (0.4 mg/mL), flash-frozen in $N_2$ (liq.) and stored at -80 °C.

*Site-specific biotinylation of VCB-AviTag using GST-BirA.*

Site-specific biotinylation of the VCB-AviTag was carried out using GST-BirA as described.[7] Briefly, the VCB-AviTag complex was first dialysed into a low salt buffer consisting of 20 mM HEPES, 20 mM sodium chloride, 1 mM TCEP, pH 7.5. The protein complex (100 µM in 952 µL of low salt buffer) was then mixed with magnesium chloride (5 µL of 1M solution) (Sigma Aldrich), adenosine triphosphate (20 µL of 100 mM solution) (Sigma Aldrich), thawed GST-BirA enzyme (20 µL of 50 µM solution) and D-Biotin (3 µL of 50 mM solution in 100% DMSO) (Sigma Aldrich) and incubated for 1 h at 30 °C in an incubator with gentle shaking (90 rpm). After this time, an additional equivalent of GST-BirA and D-biotin were added and the complex incubated for a further 1 h at 30°C. To the complex was added 100 µL of a 50% slurry of Glutathione Sepharose 4B resin (pre-equilibrated into low salt buffer) (GE Healthcare) and incubated for 30 minutes at room temperature to capture the GST-BirA. Glutathione Sepharose 4B resin and unreacted D-biotin were subsequently removed by passing the sample over a PD Minitrap G 25 desalting column (GE Healthcare) into 20 mM HEPES, 150 mM sodium chloride, pH 7.5. The extent of biotinylation was evaluated by gel-shift assay with streptavidin,[7] and found to be essentially complete. The final biotinylated complex ('VHL-biotin') was concentrated to 100 µM, snap frozen in N2 (liq.) and stored at -80 °C.

### 2. SPR binding studies

SPR experiments were performed on a Biacore T200 instrument (GE Healthcare).

**SPR binding studies (immobilised VHL).**

*Immobilization of biotinylated VHL.*

Immobilization of VHL-biotin was carried out at 25°C using either a Series S CM5 chip to which streptavidin had first been amine-coupled, or using a pre-coupled Series S SA chip. Where not expressly noted, experiments were performed using a CM5/streptavidin sensor chip. For CM5 chips, the surface was pre-equilibrated in HBS-P+ running buffer, containing 2mM tris(2-carboxyethyl) phosphine hydrochloride (TCEP), pH 7.4. Then, following activation of the surface with EDC/NHS (GE Healthcare or XANTEC) (contact time 420 sec or 600 sec, flow rate 10 µL/min), streptavidin (Sigma Aldrich) (prepared at 1 mg/mL in 10mM sodium acetate coupling buffer, pH 5.0) was immobilized by amine coupling to a density of 500-2000 RU, followed by deactivation using 1M ethanolamine. Next, the sensor chip was equilibrated in VHL running buffer consisting 20 mM 4-(2-hydroxyethyl)-1-piperazineethanesulfonic acid (HEPES), 150 mM Sodium chloride, 1 mM TCEP, 0.005% TWEEN 20, pH 7.0, 1% dimethyl sulfoxide (DMSO). Biotinylated VHL (100 nM biotinylated VHL, in VHL running buffer) was then streptavidin captured to the required surface density, using either manual injection (flow rate 10 µL/min) (CM5/streptavidin chip) or the automated wizard in the Biacore T200 control software (GE Healthcare) (SA chip, following surface preconditioning with three consecutive injections of 1M Sodium chloride in 50 mM Sodium hydroxide). For binary studies (binding of PROTAC only) the final surface density of biotinylated VHL was approximately 2000 RU; for ternary studies (binding of pre-formed PROTAC:target protein complex), multiple lower surface densities of biotinylated VHL were used (40, 80 and 120 RU) to minimise mass transfer effects. The reference surface consisted of an EDC/NHS-treated surface deactivated with 1M ethanolamine (CM5 chips) or unmodified preconditioned streptavidin surface (SA chips).

All interaction experiments (unless otherwise noted) were performed at 12°C (binary) or 25°C (ternary) in VHL running buffer. Sensorgrams were recorded at different concentrations of PROTAC (multi-cycle binary experiments) or PROTAC/target protein complex in the presence of near-saturating concentrations of target protein (single-cycle ternary experiments). For ternary experiments, the minimum concentration of target protein was selected to be approximately 20 to 50-fold in excess of the binary $K_D$ of the PROTAC/target protein interaction, to ensure a minimum binary occupancy of approximately 95 to 98%.

All ternary experiments using immobilised biotin-VHL (unless otherwise noted) were run in single-cycle kinetic mode (rather than multi-cycle mode) due to the slower dissociation kinetics of many of these complexes. This enabled reduced overall experimental run times without the need for additional surface regeneration. For binary PROTAC binding experiments using immobilised biotin-VHL the dissociation kinetics were sufficiently fast, such that these were run in multi-cycle mode.

*Binary interaction experiments (immobilised VHL).*

PROTACs (10 mM stocks in 100% DMSO) were prepared at 1 μM (300 μL) in VHL running buffer (20 mM HEPES, 150 mM Sodium chloride, 1 mM TCEP, 0.005% TWEEN 20, pH 7.0) containing 1% DMSO. This stock solution was then serially diluted in VHL running buffer containing final 1% DMSO (either 11-point two-fold serial dilution, 1000 nM - 0.98 nM final concentration of PROTAC, 150 μL sample volume; or 5-point five-fold serial dilution, 1000 nM - 1.6 nM final concentration of PROTAC; 240 μL sample volume). Solutions were injected individually (duplicate wells) in multi-cycle kinetic format without regeneration (contact time 210 sec, flow rate 50 μL/min, dissociation time 300 sec) using a stabilisation period of 30 sec and syringe wash (50% DMSO) between injections.

*Ternary interaction experiments (immobilised VHL).*

PROTACs (10 mM in 100% DMSO) were initially prepared at 1 μM or 200 nM in VHL running buffer with a concentration of 2% DMSO. This solution was mixed 1:1 with a solution of 50 μM of the corresponding bromodomain target protein in VHL running buffer without DMSO, to prepare a final solution (300 μL) of 500nM or 100nM PROTAC and 25 μM bromodomain in VHL running buffer containing 1% DMSO. This complex was then serially diluted in VHL running buffer containing 2 μM bromodomain and 1% DMSO (5-point five-fold serial dilution, 500 nM – 800 pM or 100 nM – 160 pM final concentration of PROTAC, 25 μM – 2 μM final concentration of bromodomain). For ternary experiments, solutions were injected sequentially in single-cycle kinetic format without regeneration (four replicate series per experimental repeat, contact time 100 sec, flow rate 100 μL/min, dissociation time 720 sec) using a stabilisation period of 30 sec and syringe wash (50% DMSO) between injections. High flow rates and multiple surface densities were used to minimise mass transfer effects. At least two series of blank injections (VHL running buffer containing 2 μM bromodomain and 1% DMSO) were performed for all single cycle experiments to be used for blank subtraction.

*Ternary interaction experiments varying the PROTAC:target ratio (MZ1:Brd4$^{BD2}$) (immobilised VHL)*

These were run similarly to other ternary interaction experiments, with the modifications that these experiments were recorded in multi-cycle format at 12°C using a pre-coupled Series S SA chip to which approximately 500 RU of biotin-VHL had been immobilised.

MZ1 (10 mM stock in 100% DMSO) was serially diluted to prepare a 5-point concentration series (2 µM, 400 nM, 80 nM, 16 nM, or DMSO vehicle) in VHL running buffer with a final concentration of 2% DMSO. This solution was mixed 1:1 in a plate format with a corresponding 5-point concentration series of Brd4$^{BD2}$ (2 µM, 400 nM, 80 nM, 16 nM, or running buffer vehicle) in VHL running buffer without DMSO, to ultimately prepare final solutions (300 µL) of PROTAC:BD in varying ratios (1:1, 1:5, 1:25, 1:125; as depicted in Figure S5(a)(i)) in VHL running buffer containing 1% DMSO. Solutions were injected sequentially in multi-cycle kinetic format without regeneration (single injection per concentration, contact time 120 sec, flow rate 80 µL/min, dissociation time variable, 2500 to 4000 sec) using a stabilisation period of 30 sec and syringe wash (50% DMSO) between injections. A single blank injection (DMSO vehicle) at each concentration of Brd4$^{BD2}$ was measured and used for background subtraction of all sensorgrams with the same Brd4$^{BD2}$ concentration.

**Reversed-format SPR binding studies (immobilised Brd4$^{BD2}$)**

*Immobilization of biotinylated Brd4$^{BD2}$.*

Immobilization of biotinylated Brd4$^{BD2}$ was carried out at 25°C using a pre-coupled Series S SA chip. The sensor chip was equilibrated in VHL running buffer consisting 20 mM 4-(2-hydroxyethyl)-1-piperazineethanesulfonic acid (HEPES), 150 mM Sodium chloride, 1 mM TCEP, 0.005% TWEEN 20, pH 7.0, 1% dimethyl sulfoxide (DMSO). Biotinylated Brd4$^{BD2}$ (100 nM biotinylated Brd4$^{BD2}$, in VHL running buffer) was then streptavidin captured to the required surface density using the automated wizard in the Biacore T200 control software (GE Healthcare), following surface preconditioning with three consecutive injections of 1M sodium chloride in 50 mM sodium hydroxide. For binary studies (binding of PROTAC only) the final surface density of biotinylated Brd4$^{BD2}$ was approximately 2800 RU; for ternary studies (binding of pre-formed PROTAC:target protein complex), the final surface density of biotinylated Brd4$^{BD2}$ was ~400 RU. The reference surface consisted of an unmodified preconditioned streptavidin surface. Interaction experiments were performed at 12°C in VHL running buffer, as described for immobilised VHL.

*Preliminary binary interaction experiments (immobilised Brd4$^{BD2}$).*

PROTACs (10 mM stocks in 100% DMSO) were prepared at 250 nM (300 µL) in VHL running buffer (20 mM HEPES, 150 mM Sodium chloride, 1 mM TCEP, 0.005% TWEEN 20, pH 7.0) containing 1% DMSO. This stock solution was then serially diluted in VHL running buffer containing final 1% DMSO (either 5-point five-fold serial dilution, 250 nM – 0.4 nM final concentration of PROTAC; 240 µL sample volume). Solutions were injected individually (duplicate wells) in single-cycle kinetic format without regeneration (contact time 100 sec, flow rate 100 µL/min, dissociation time variable 900 - 3000 sec) using a stabilisation period of 30 sec and syringe wash (50% DMSO) between injections. In the case of MZP61 (binary), a regeneration step was used consisting of a single injection of 2 μM VHL in running buffer containing 1% DMSO (contact time 100 sec, flow rate 100 μL/min, dissociation time 900 sec), which caused rapid dissociation of the formed ternary complex. Two series of blank injections were performed for all single cycle experiments.

*Preliminary ternary interaction experiments (immobilised Brd4$^{BD2}$).*

PROTACs (10 mM in 100% DMSO) were initially prepared at 500 nM in VHL running buffer with a concentration of 2% DMSO. This solution was mixed 1:1 with a solution of 50 μM of VHL target protein in VHL running buffer without DMSO, to prepare a final solution (300 μL) of 250nM PROTAC and 25 μM VHL in VHL running buffer containing 1% DMSO. This complex was then serially diluted in VHL running buffer containing 2 μM VHL and 1% DMSO (5-point five-fold serial dilution, 250 nM – 400 pM final concentration of PROTAC, 25 μM – 2 μM final concentration of VHL). For preliminary ternary experiments, solutions were injected sequentially in single-cycle kinetic format without regeneration (one series per experimental repeat, contact time 100 sec, flow rate 100 μL/min, dissociation time variable 900 - 3000 sec) using a stabilisation period of 30 sec and syringe wash (50% DMSO) between injections. Two series of blank injections were performed for all single cycle experiments.

**SPR data analysis.**

Data were analysed using either or Scrubber (BioLogic Software) or Biacore T200 Evaluation Software (GE Healthcare). Sensorgrams from reference surfaces and blank injections were subtracted from the raw data (double-referencing) and the data was solvent-corrected prior to analysis. To calculate the association rate ($k_{on}$), dissociation rate ($k_{off}$), and dissociation constant ($K_D$), data from all binary (multi-cycle) and ternary (single cycle) experiments were fitted using a 1:1 Langmuir interaction model, with a term for mass-transport included. For experiments conducted at multiple surface densities, this was performed using global fitting of data from all surface densities simultaneously (zip fitting in Scrubber) (BioLogic Software) (as described in Figure S3). For some ternary experiments, nonspecific effects were observed during the association phase at the top concentration only; in these cases, the fifth injection only was not included in global fitting.

### 3. Fluorescence polarization (FP) cooperativity assay (VHL binding)

All FP measurements were taken using a PHERAstar FS (BMG LABTECH) with fluorescence excitation and emission wavelengths (λ) of 485 nm and 520 nm, respectively. FP competitive binding assays were performed in triplicate on 384-well plates (#3575, Corning) with a total well volume of 15 μL. Each well solution contained 10 nM of FAM-labelled HIF-1α peptide (FAM-DEALAHypYIPMDDDFQLRSF, $K_D$ = 3 nM as measured by a direct FP titration), 15 nM of VCB protein, and decreasing concentrations of MZ1 (14-point serial 2-fold dilutions starting from 50 μM) or MZ1:bromodomain (14-point serial 2-fold dilution starting from 10 μM MZ1:100 μM bromodomain into buffer containing 1 μM bromodomain) in 100 mM Bis-tris propane, 100 mM sodium chloride, 1 mM TCEP, pH 7, with a final DMSO concentration of 1%. To obtain percentage of displacement, control wells containing peptide in the absence of protein (maximum displacement), or VCB and peptide with no compound (zero displacement) were also included. These values were then fitted by nonlinear regression using Prism (GraphPad, version 7.03) to determine average $IC_{50}$ values and standard error of the mean (SEM) for each titration. A displacement binding model was used to back-calculate inhibition constants ($K_I$) from the measured $IC_{50}$ values, as described previously.[8]

### 4. Cell biology and degradation studies

**Cell lines and culture.**

HEK293 cells (ATCC) were grown in DMEM (Invitrogen) supplemented with 10% v/v fetal bovine serum (Brazil origin, Life Science Production) at 37 °C and 5% $CO_2$ in a humidified atmosphere. Cells were split 1-2 times per week when 90% confluent and were not used beyond passage 25. Cells were routinely checked for mycoplasma contamination using Mycoalert detection kit (Lonza).

**Degradation time course assays.**

HEK293 cells were seeded at 4-8 x $10^5$ cells/well of 6 well plates 12-24 h before treatment. Cells were treated with 333 nM MZ1 or 0.1% v/v DMSO and lysed over a range of time points up to 7 h after treatment. Upon lysis, cells were washed twice in ice cold PBS (Invitrogen) then lysed and scraped in 80 μL/well of ice-cold lysis buffer containing 50 mM Tris hydrochloride pH 7.4, 150 mM sodium chloride, 1 mM EDTA pH 7.4, 1 % v/v Triton X-100, protease inhibitor cocktail (Roche). Lysates were sonicated, cleared by centrifugation at 4 °C, at 15800 x *g* for 10 mins and the supernatants stored at -80 °C. Protein concentration was determined by BCA assay (Pierce) and the absorbance at 582 nm measured by spectrophotometry (NanoDrop ND1000). Samples were run on SDS-PAGE using NuPAGE Novex 4-12% Bis-Tris gels (Invitrogen) with 40 μg total protein/well, transferred to nitrocellulose membrane and blocked with 3 % w/v BSA (Sigma) in 0.1% TBST. Blots were incubated in anti-Brd2 (1:2000, abcam #ab139690), anti-Brd3 (1:500, abcam #ab50818), anti-Brd4 (1:1000, abcam #ab128874), or anti-β-actin (1:2500, CTS #4970S) antibody overnight at 4 °C with rotation. Blots were then incubated in goat anti-mouse or donkey anti-rabbit IRDye 800CW secondary antibodies (1:10,000, LICOR #925-32210 and #926-32213) for 1 h at room temperature with rotation. Bands were detected using a ChemiDoc (BioRad) and quantified (Image Studio Lite, version 5.2) with normalisation to β-actin and DMSO control per time point. Data are the average of two Western blots per protein from three biological repeats.

Degradation data were plotted and fitted by nonlinear regression over the initial degradation period (from experiment start until the point of maximum degradation, $D_{max}$) using a single-phase exponential decay model in Prism (GraphPad, version 7.03) to obtain estimates for degradation rate ($\lambda$), y-intercept ($y_0$) and plateau.[9] Degradation half–life ($t_{1/2}$) was calculated from the fitted degradation rate.
